# Inferring single-cell transcriptomic dynamics with structured latent gene expression dynamics

**DOI:** 10.1101/2022.08.22.504858

**Authors:** Spencer Farrell, Madhav Mani, Sidhartha Goyal

## Abstract

Gene expression dynamics provide directional information for trajectory inference from single-cell RNA-sequencing data. Traditional approaches compute local RNA velocity using strict assumptions about the equations describing transcription and splicing of RNA. Not surprisingly, these approaches fail where these assumptions are violated, such as in multiple lineages with distinct gene dynamics or time-dependent kinetic rates of transcription and splicing. In this work we present “LatentVelo”, a novel approach to compute a low-dimensional representation of gene dynamics with deep learning. Our approach embeds cells into a latent space with a variational auto-encoder, and describes differentiation dynamics on this latent space with neural ordinary differential equations. These more general dynamics enable accurate trajectory inference, and the latent space approach enables the generation of a latent “dynamics-based” embedding of cell states. To model multiple distinct lineages, LatentVelo infers a latent regulatory state that controls the dynamics of an individual cell. With these lineage-specific dynamics LatentVelo can predict latent trajectories, describing global inferred developmental path for individual cells, rather than just outputting local RNA velocity vectors. The dynamics-based embedding also enables concurrent batch correction of cell states and RNA velocity, outperforming comparable auto-encoder based batch correction methods that do not consider gene expression dynamics. Finally, the flexible structure of LatentVelo enables additional of new regulatory constraints required to integrate multiomic data. LatentVelo is available at https://github.com/Spencerfar/LatentVelo.

## I. INTRODUCTION

Single-cell RNA sequencing enables the inference of developmental trajectories with methods that computationally reconstruct the developmental process. Traditional approaches are based on the similarity of static snapshots of RNA for individual cells [1–3]. RNA velocity extends these approaches by modelling the dynamical relationship between newer unspliced RNA and mature spliced RNA to infer the direction of the dynamics from these static snapshots. Recent techniques have also been developed to analyze RNA velocity, aiming to improve and expand upon traditional trajectory reconstruction methods [4–9].

However, RNA velocity methods have been limited by a reliance on strict modelling assumptions. In the original formulation, Velocyto [10] modelled cells at an assumed steady-state, with a linear relationship between unspliced and spliced RNA. scVelo [11] relaxed the steady-state assumption by assuming a strict form for transcription rates, and modelled time-dependent transient cell-states by fitting a set of linear differential equations, treating the unobserved developmental time as a latent variable inferred per cell and per gene. These methods encounter problems with complex dynamical features such as a transcriptional boost, lineage-dependent kinetics, and weak unspliced signal [12–14].

We have developed “LatentVelo”, an approach to address these problems in RNA velocity estimation. Our key insight and distinguishing feature is that LatentVelo embeds cell states in a learned latent space, and infers structured dynamics on this latent space that incorporate the causal structure of RNA velocity dynamics, rather than learn RNA velocity on gene-space with strict linear dynamics. Since the latent dynamics and embedding are learned together, the latent embedding of cell states is informed by the dynamics. Lineage or time-dependent dynamics are enabled by modelling state-dependent regulation of transcription with a latent regulatory state. By learning dynamics in a latent space, LatentVelo can extract the low-dimensional dynamics describing cell differentiation. Additionally, modelling the latent space dynamics enables batch correction and the incorporation of additional information such as annotated cell-type labels as in scANVI [15] or temporal information from sequencing batches. LatentVelo also enables constructing general dynamical models, and enabling various structured models of regulation and multi-omic data.

Recently, several other methods have been proposed to address some of these issues to improve the accuracy and reliability of RNA velocity methods. VeloAE [16] develops an extension of the steady-state model by projecting high-dimensional noisy unspliced and spliced vectors onto a lower dimension latent space with an autoencoder, and then uses the steady-state linear model to estimate velocities on this latent space. UniTVelo [17] allows for a more general form of spliced RNA dynamics which relaxes assumptions on the transcription rate, includes a unified cell-dependent latent time (rather than per gene as in scVelo), and introduces a method of dealing with low signal-to noise in unspliced reads. DeepVelo [18] uses a deep neural network to estimate cell-specific kinetic rate parameters to model variable and lineage-dependent kinetics. VeloVAE [19] uses a variational Bayesian approach to model a unified cell-dependent latent time, cell-specific transcription rates, and celltype specific splicing and degradation rates. MultiVelo [20] integrates chromatin accessibility into the linear dynamical model to improve velocity estimates. Alternatively, work has been done on gene selection for velocity analysis [12, 21].

LatentVelo has similarities with these methods. LatentVelo embeds cell states in a latent space with an auto-encoder like VeloAE, but includes dynamics rather than just modelling steady-state cells. LatentVelo models dynamics with a unified cell developmental time like UniTVelo and VeloVAE. Like DeepVelo and VeloVAE, LatentVelo models lineage-dependent dynamics. LatentVelo also includes the scTour model [22] as a special case, where only the dynamics of a single data-type are considered (e.g. just spliced RNA).

We benchmark LatentVelo with synthetic data and real developmental, regeneration, and reprogramming data. We show that our approach significantly outperforms the traditional linear RNA velocity methods for inferring transitions between cell-types. We also show that our approach is able to do batch correction of RNA velocity by projecting batches onto a common latent space independent of technical variation between the samples. Batch correction is unaddressed by other models of RNA velocity, presenting a novel application of LatentVelo. Additionally, rather than just output RNA velocity vectors, LatentVelo infers developmental trajectories for individual cells, describing the inferred path a specific cell took to reach its current state. This is distinct from the traditional velocity methods.

## II. OVERVIEW OF LATENTVELO

### A. RNA velocity

RNA velocity methods estimate the direction of differentiation using static snapshots of unspliced and spliced RNA. The quantity of unspliced and spliced RNA are determined by the processes of processes of transcription, splicing, and degradation of RNA:

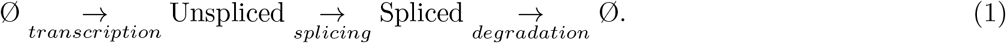

The objective is to overcome the limitations of snapshot scRNAseq observations by using a model that imposes the causal relationship of splicing, linking observed quantities of unspliced and spliced RNA. The goal is to estimate the time-derivative of the spliced RNA for single-cells 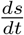 in terms of the amount unspliced and spliced RNA, which characterizes the future direction of differentiation for a cell.

The traditional model for RNA velocity uses linear rate equations for the transcription, splicing, and degradation dynamics per gene *g*,

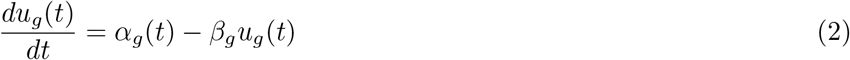

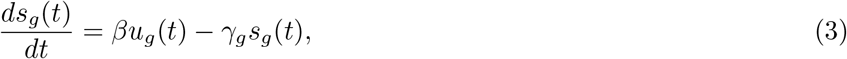

where *t* is an unknown developmental time that is either eliminated in the steady-state model (Velocyto) [10], or inferred per gene and per cell in the linear dynamical model (scVelo) [11]. Instead of assuming the accuracy of these linear models in accounting for the data, we construct a general deep-learning model that incorporates only the causal relationships outlined in (1), while remaining agnostic to the underlying kinetics of the process. The goal is to encode the causal relationship, but generalize the linearity assumption. As such the approach is more data-driven.

### B. LatentVelo

LatentVelo is formulated as a variational auto-encoder (VAE) that embeds RNA count data into latent states 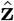 along with a latent developmental time 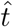 per cell, which provides a time-ordering of the cells. Variational auto-encoders are approximate Bayesian models that use a neural network based encoder to approximate the posterior distribution of a model [23, 24]. In our case, we estimate the posterior distribution 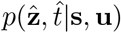 of latent states and latent developmental times.

Dynamics on this latent space are described by a neural ordinary differential equation [25], using the inferred latent time as the time variable in the differential equation. The basic form of the latent dynamics are,

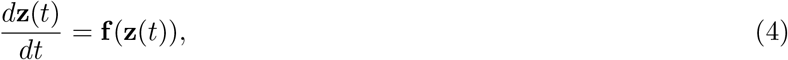

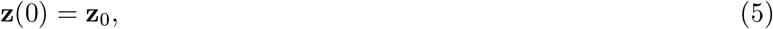

where the dynamics are determined by a neural network **f** and an initial state **z**_0_. Note that the initial state **z**_0_ is *learned* together with the dynamics and autoencoder, in contrast to pseudotime methods which specify a root cell. These dynamics are incorporated into the VAE by constraining the encoded state state 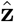 with the solution to these dynamics at the corresponding latent time for the cell 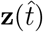, e.g. minimizing 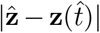. This encourages the encoder to output latent states 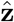 that follow the latent dynamics, and encourages the latent dynamics **z**(*t*) to match the output of the encoder. The result is that LatentVelo learns a latent embedding that is informed by the dynamics.

These dynamics are considered “unstructured” in the sense that **f** is a general dense feed-forward neural network, and interactions between all components of **z** are allowed. A very similar model with these unstructured dynamics was also recently developed in scTour [22]. However, these unstructured models do not utilize biological knowledge of the dynamics of splicing. We add this structure by separately representing the different data modalities in the latent space by decomposing **z** = (**z**_*u*_, **z**_*s*_) and **f** = (**f**_*u*_, **f**_*s*_),

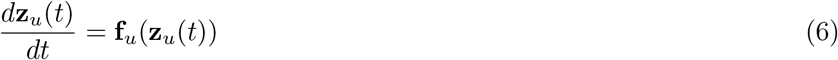

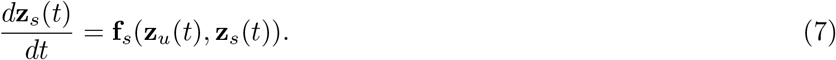

The structure comes in by separately modelling spliced **z**_*s*_ and unspliced **z**_*u*_ latent representations, and restricting the interactions between them to model splicing dynamics. This results in an estimate of a latent representation of RNA velocity, *d***z**_*s*_*/dt*. This model is a generalized form of traditional RNA velocity methods – modelling the dynamics of transcription, splicing, and degradation with non-linear functions of spliced and unspliced RNA, and including interactions between different genes through this low-dimensional latent space.

However, this form of dynamics does not address the issue of multiple lineages. Since the dynamics start from a single initial state **z**_0_ and all cells follow the same ODEs, there is only one unique solution – which means no branching for different lineages. Lineage-dependent dynamics can be incorporated by including regulation of transcription with state-dependent dynamics of a new latent variable controlling regulation **z**_*r*_,

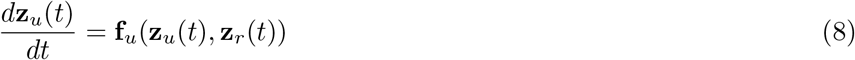

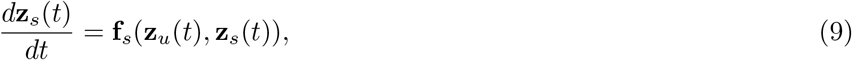

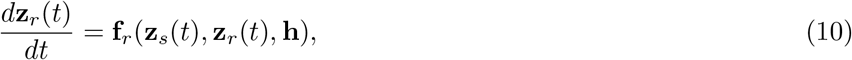

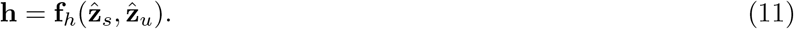

The variable **h** is a constant state-dependent parameter controlling the regulatory dynamics for a particular cell. Note that **h** depends on the observed cell state because it depends on the encoded 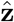, *not* the trajectory **z**(t). Since h is estimated from the observed cell-state, it enables dynamics to depend on cell-states or lineages. Since the dynamics are deterministic and start from a constant **z**(0), without **h** we would expect no branching of the dynamics. We can think of **h** as conditioning the dynamics to follow a particular branch of the system. This enables us to model complex multi-lineage dynamics. Note that while the VAE includes stochasticity in the latent embedding, there is no stochasticity in the dynamics. We address this in the discussion as a future direction.

We further enforce the direction of splicing (unspliced → spliced) by regularizing a positive correlation between 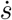 and *u* and a negative correlation between 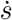 and *s*, e.g., 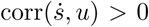 and 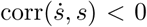 per gene. By only weakly including this regulation, we allow time-dependent rates rather than the strict linear assumption of some previous models. A similar form of regularization is also done in DeepVelo [18]. Further details are in the Methods section V B.

Figure 1 visualizes the structure of LatentVelo described in Equations 8 to 11 in a), and demonstrates some of the features of LatentVelo on a synthetic dataset. The UMAP representation of this dataset is shown in b), highlighting the direction of differentiation with increasing simulation time, and specific milestone cell clusters. In c) we show the latent time and velocities inferred by LatentVelo. LatentVelo’s latent time is similar to a pseudotime, and velocities are a low-dimensional representation of RNA velocity. In d) we show UMAP representation of the spliced latent state, which is a latent embedding of cell states informed by the transcriptomic dynamics. In e) we plot the latent regulatory state *z*_*r*_ vs latent time, showing that LatentVelo infers distinct dynamics on the two branches.

**FIG. 1.**
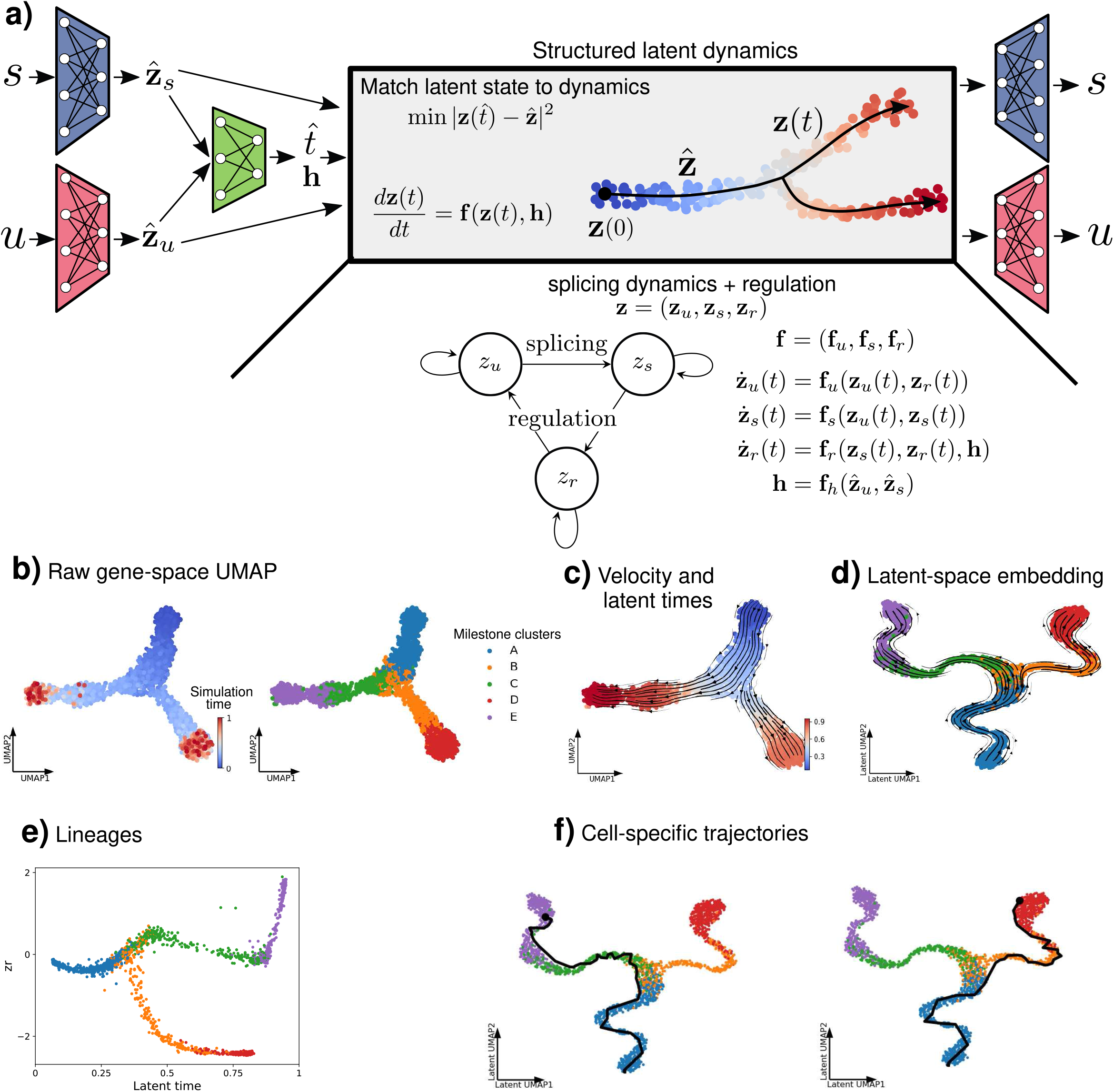
LatentVelo. **a)** LatentVelo is a variational autoencoder with structured dynamics on the latent space. An encoder (left trapezoids) encodes the transcriptomic cell state (i.e. unspliced (*u*) and spliced (*s*) RNA counts) into corresponding latent states 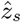 and 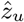, and a latent developmental time *t* and regulation state **h**. The regulation state h conditions dynamics for each cell to follow a particular branch. We match these latent states to the dynamics obtained by structured dynamics on this latent space, **z**(*t*), described by ODEs starting from an initial state **z**(0). These structured dynamics can take a variety of forms, here we show regulated splicing dynamics incorporating state-dependent regulation of transcription. In the shown notation, we use 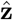 to denote latent cell states estimated directly from the encoder with the input data, and **z**(*t*) to represent latent cell states estimated from the latent dynamics. When fitting the model, the distance between the encoder latent states and the dynamics latent states is minimized. **b)-f)** We demonstrate LatentVelo on an example synthetic dataset. In **b)**, we show the gene-space UMAP for this dataset, highlighting the developmental trajectory with the simulation time and celltype transitions with the milestone clusters. We show LatentVelo’s ability to infer velocity and latent time **c)** and a latent embedding **d)**. We show that **z**_r_ infers branching in **e)**, and show that we can infer cell-specific trajectories in **f)**, with trajectories for two cells on different branches of the bifurcation.

Current RNA velocity methods estimate the spliced velocity of each observed cell. In Figure 1f), we show that LatentVelo infers cell-specific trajectories on the latent space, instead of simply inferring local cell velocities. Typically, RNA velocity has been interpreted by visualizing global velocity fields of smoothed local cell velocities projected onto a 2-dimensional representation representation of the data (e.g. UMAP or tSNE). LatentVelo’s inferred trajectories offer a new way of interpreting RNA velocity, without resorting to the 2-dimensional representation of velocities. These trajectories directly indicate the path of differentiation, which more clearly indicates transitional celltypes.

LatentVelo is not limited to the specific structure of transcription and splicing used here. Any desired structure to the latent dynamics can be implemented, for example regulation of splicing can be included with an interaction from *z*_*r*_ to *z*_*s*_. This generic nature of the model enables modeling multi-omic dynamics, by specifying a particular structure to the dynamics linking the different data modalities. For example in the Supplemental Information, we demonstrate an example where we model both transcription and splicing from RNA sequencing together with chromatin accessibility from ATAC-seq. We have not explored other variations to the splicing and regulation structure.

The standard VAE version of the model uses a standard Gaussian prior on the latent state 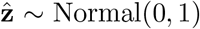 and a logit-normal prior on the latent times 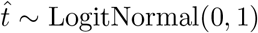. We can add more structure to the latent embedding by including cell-type information, similar to scANVI [15]. This approach modifies the prior of the latent space to be cell-type specific, rather than a standard Gaussian for all cells. This modification prevents the prior from ignoring biologically relevant clusters, and in particular, we show below that this improves batch correction. The full details of this modification are in the Methods section V C.

Since LatentVelo has many free parameters, it can be useful to include the experimental time of observation of cells to further constrain the dynamics. We do this by adding the correlation between the inferred latent time and experimental time to the loss function. The details of this are in Methods section V D.

### C. Evaluation of LatentVelo

We take a quantitative approach to evaluate RNA velocity by comparing with ground truth velocity on synthetic data, and the direction of known cell-type transitions in real data.

To evaluate LatentVelo on synthetic data, we use the cosine similarity between ground truth and estimated velocities. Since many genes are not directly involved in the differentiation process, we compute this metric on lower dimensional embeddings; the model latent space and 50-dimensional principle component (PC) space.

For real data, we use datasets where the direction of differentiation is known. To quantitatively evaluate models, we use a modified version of the Cross-Boundary Directedness score used for UniTVelo and VeloAE [16, 17]. This score measures the velocity direction of cells on the boundary between two cell-types.

We make two modifications of this score. In previous work this score was computed on a 2D UMAP embedding, but we found this sometimes resulted in inconsistent values not seen in higher-dimensional embeddings. Therefore, we use multiple higher-dimensional embeddings (model latent space, higher dimensional PCA). Additionally, we modify the score to be more robust to noisy boundaries between cell-types by only checking that the direction of the velocity of a cell is directed towards any cell of the expected cell-type, rather than any specific cell (see Methods section V F).

Our updated version quantifies the probability that a cell is likely to transition in the specified direction. Therefore, we can interpret scores above 0.5 to indicate a likely transition in that direction. Note that since this score depends on well-defined boundaries, noisy boundaries can lower the score.

We also use the Inner-Cluster Coherence score used for UniTVelo and VeloAE [16, 17]. This score evaluates the coherence of velocity direction for neighboring cells within a cluster or cell-type. At a maximum value of 1, neighboring cells have the same velocity direction. A similar consistency score was also used with scVelo [11] and DeepVelo [18].

We use the kBET and iLISI batch correction metrics and the cLISI biological cluster conservation metrics to evaluate our models performance for batch correction [26]. kBET and iLISI measure how well batches are integrated together, and cLISI measures how well biological clusters are retained when batch correcting. These metrics only evaluate batch correction of gene expression. To evaluate the batch correction of RNA velocity, we compute the average cosine similarity between neighboring cells.

## III. RESULTS

### A. LatentVelo infers cell fate trajectories

In Figure 2, we demonstrate LatentVelo on a pancreatic endocrinogenesis dataset showing differentiation of endocrine progenitors into 4 terminal states: Alpha, Beta, Delta, and Epsilon cells [27]. In 2a), a UMAP plot of annotated celltypes in this dataset are shown. In b) we show the latent time and velocities estimated by LatentVelo on this UMAP plot, which show the differentiation of progenitors into the terminal cell states. In c), we visualize the 2-dimensional latent regulatory state **z**_*r*_, which controls the dynamics of each cell. The 4 terminal celltypes cluster into separate regions, showing that LatentVelo learns distinct dynamics for these 4 terminal states. In contrast, traditional RNA velocity methods learn the same parameters for all celltypes.

**FIG. 2.**
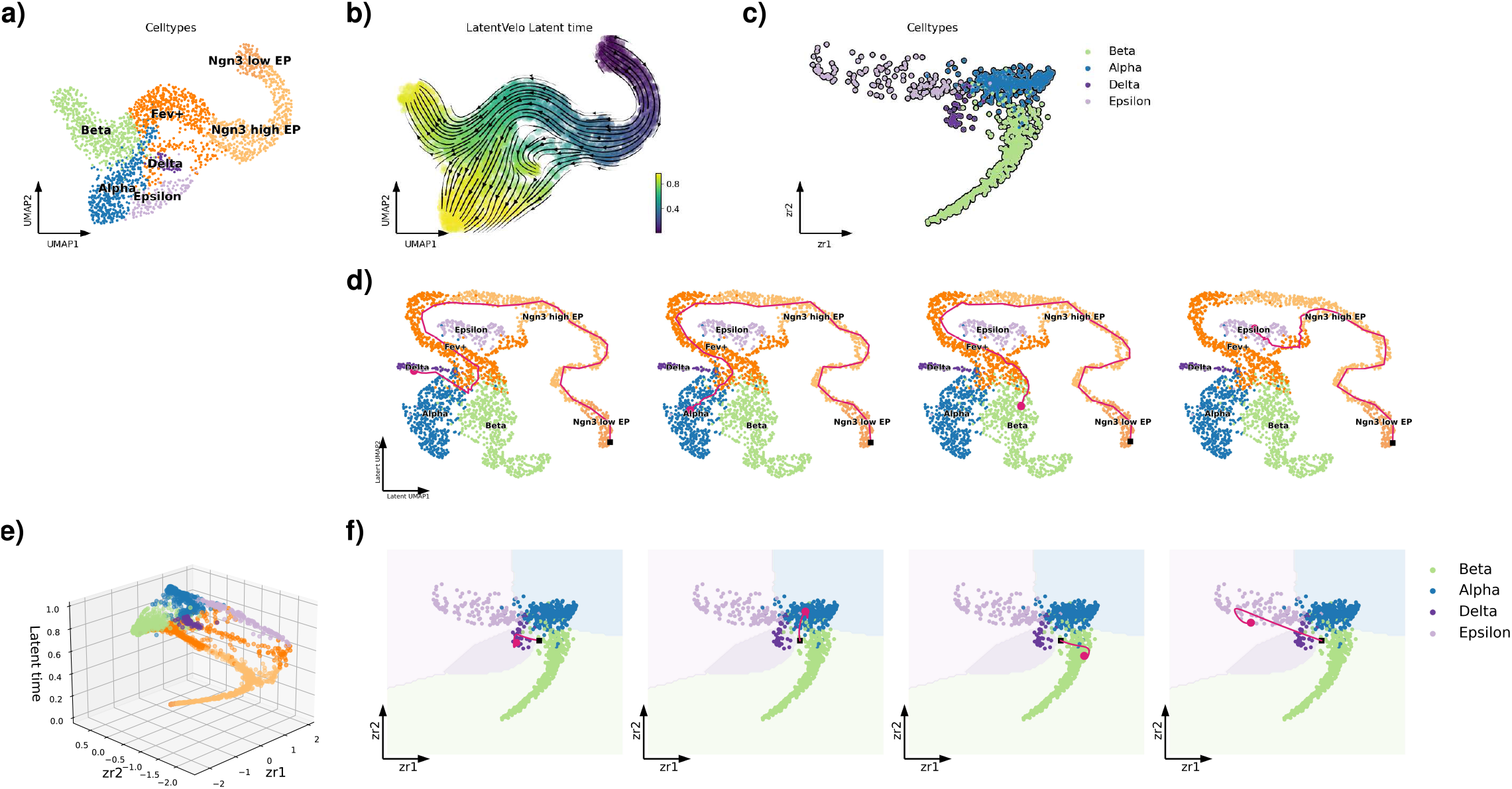
LatentVelo infers latent cell dynamics. **a)** UMAP plot showing the differentiation of endocrine progenitors into 4 terminal states, Alpha, Beta, Delta, and Epsilon cells. **b)** We show the velocity and latent time inferred by LatentVelo, indicating the direction of differentiation. **c)** We also show the inferred latent regulatory states in LatentVelo, highlighting the distinct dynamics for the terminal states. **d)** We show the latent trajectories inferred by LatentVelo for 4 different cells, corresponding to a Delta, Epsilon, Beta, or Alpha cell. Plot is shown for a UMAP representation of the spliced latent state. **e)** Latent regulatory state dynamics, showing a 3-dimensional plot of *z*_*r*1_ vs *z*_*r*2_ vs latent time. **f)** We show a scatter plot of the terminal states for *z*_*r*1_ vs *z*_*r*2_. Dynamics start from the initial state z_0_ at the black square (the *learned* initial state) and evolve with the estimated *h* and follow the magenta path until terminating at the magenta circles at the estimated 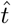.

In 2d) we choose 4 cells, each belonging to one of the 4 terminal states, and plot the inferred dynamics for these cells. The black square denotes the initial cell state **z**_0_ at time 0, which is *learned* together with the dynamics and autoencoder, and the magenta circle is the final state of the trajectory at 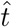. In each of these trajectories, the encoder of the model infers the conditioning variable **h**, which determines the dynamics of **z**_r_ which regulates the dynamics of a given cell to fall along a particular lineage. These trajectories more clearly delineate the differentiation of the 4 terminal celltypes in comparison to 2b). In particular, the development of epsilon cells is separated from Alpha cells and is more cleanly seen as a separate terminal state.

In 2e), we show the dynamics of the latent regulatory state **z**_*r*_, where both dimensions are shown against latent time in a 3-dimensional plot. In f) we show trajectories of **z**_*r*_ on a 2-dimensional plot only showing the terminal states. **z**_*r*_ evolves from the initial black square to the respective regions representing the 4 terminal cell types. This plot visualizes the branching dynamics for these celltypes.

A second example showing the differentiation of human hematopoietic stem cells into 6 different celltypes is shown in Supplemental Figure S1. These trajectories suggest transitions that are unclear from simply plotting a velocity field on 2-dimensional tSNE representation of the data.

LatentVelo is similar yet distinct in this aspect of inferring trajectories to recent approaches that utilize RNA velocity dynamics in further analyses. CellRank [4] uses a cell to cell transition matrix, which can be constructed with RNA velocity, to extract initial states, terminal states, and absorption probabilities of cells transitioning to a particular terminal state. The difference with LatentVelo is that we infer the past trajectory, rather than the probability of a future particular fate. Dynamo [5] uses RNA velocity estimates to interpolate a smooth velocity field, from which various differential geometric quantities can be extracted, and can the velocity field can be used to simulate trajectories for a specific initial starting state.

Inferring these trajectories is more difficult than simply estimating velocity direction. In Supplemental Figure S4, we show on an intestinal organoid dataset [28] that estimated velocities can show expected celltype transitions (visually on UMAP and quantitatively with CBDir) while inferred trajectories are poor – the final state of the trajectory 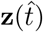 does not match the directly encoded state from the data 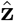. We also show that this can be improved by increasing the dimensions of the latent space to include more information. This procedure can generally be used to determine the required dimension of the latent space – we can increase the dimension until trajectories terminate close to the directly encoded state from the encoder.

### B. LatentVelo infers early separation of trajectories for reprogrammed and dead-end cells

To demonstrate LatentVelo’s ability to learn biologically meaningful features in its latent representation, we use a dataset of mouse embryonic fibroblasts (MEF) reprogramming toward induced endoderm progenitors (iEP) [29]. In this experiment, over-expression of key transcription factors drives fibroblasts to potentially undergo reprogramming, or develop into a dead-end state. Celltagging is used to identify longitudinal trajectories of reprogramming. On this dataset, scVelo incorrectly shows differentiation from reprogrammed to dead-end cells, and shows no route to reprogramming [4].

In Figure 3 a)-e), we show a tSNE plot of the data. We show the velocity estimated by scVelo and LatentVelo overlayed with their respective inferred latent times. We also show colors showing experimental time course, LatentVelo regulatory parameter *z*_*r*_, and reprogramming outcome (purple dead-end, green reprogrammed). LatentVelo estimates latent time that correlates with the experimental time-course, and velocity that indicates transitions to both dead-end and reprogrammed cells. In contrast, scVelo shows differentiation of reprogrammed cells towards dead-end cells.

**FIG. 3.**
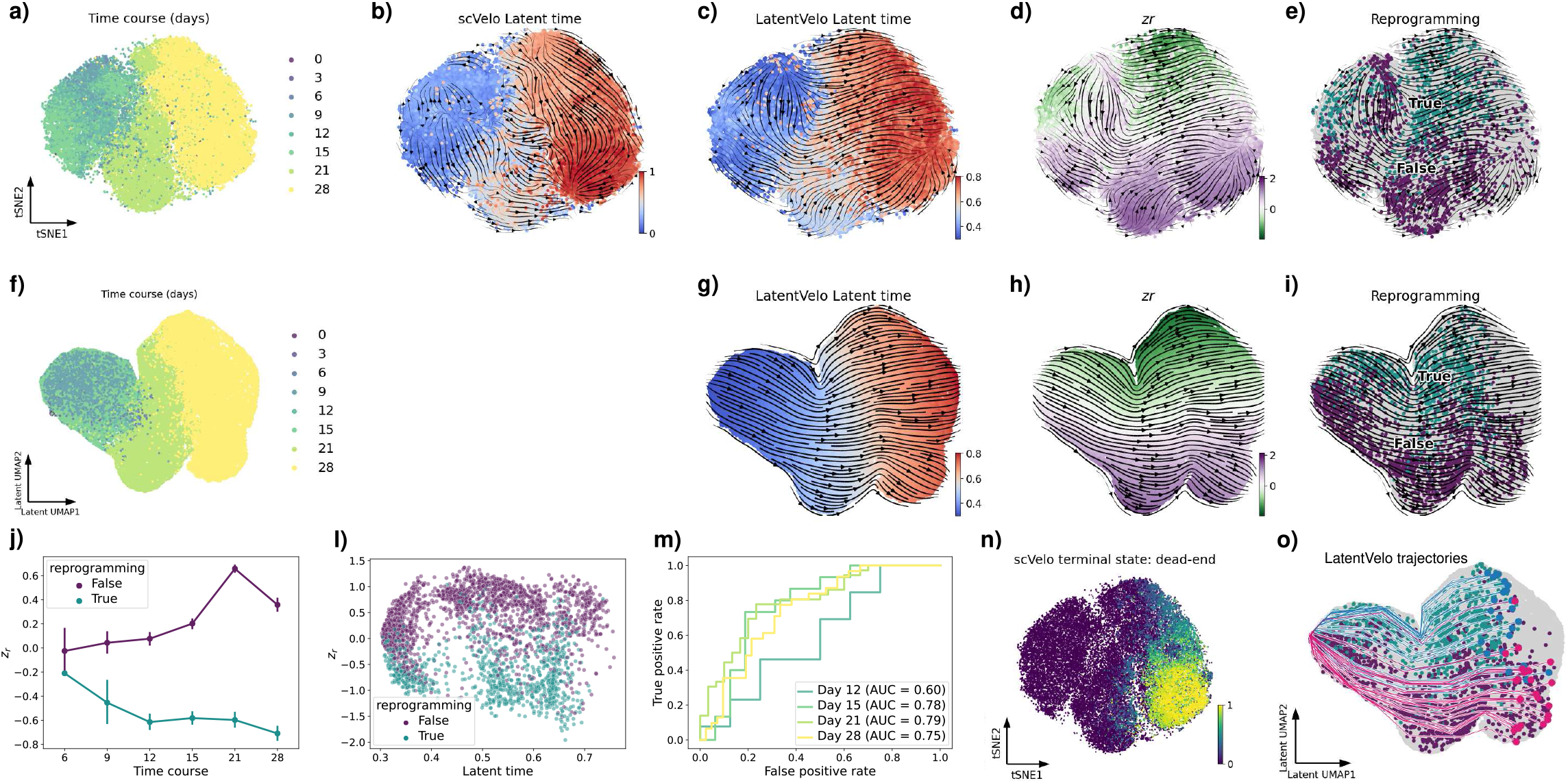
Reprogramming dynamics of mouse fibroblasts. **a)-e)** t-SNE representation of mouse embryonic reprogramming data [29]. We color this plot by experiment time course in a). We plot scVelo velocity and latent time and LatentVelo velocity and latent time in **b)** and **c)**. In **d)** and **e)** we show LatentVelo velocity with regulatory parameter *z*_*r*_ and reprogramming outcome (turquoise reprogrammed, purple dead-end) from the experiment. We see that scVelo incorrectly predicts differentiation from reprogrammed cells to dead-end cells, whereas LatentVelo shows no such transitions and shows a route to reprogramming from early times. The regulation parameter *z*_*r*_ in LatentVelo is highly correlated with reprogramming, and describes distinct dynamics for different values of *z*_*r*_ corresponding to reprogramming and dead-end cells. **f)-i)** We compute a UMAP representation of the latent spliced latent state **z**_*s*_, and color by the variables in the first row. On this learned latent space, dynamics are clearly separated by *z*_*r*_, corresponding to the reprogramming outcome. **j)-o)** We show that reprogramming outcome is strongly correlated with *z*_*r*_, by showing experimental time course and inferred latent time vs *z*_*r*_, colored by reprogramming outcome. In m) we include a ROC curve from a one-dimensional logistic regression showing *z*_*r*_ is predictive of reprogramming outcome. In n) we also show the original tSNE embedding colored by the predicted fate probabilities by CellRank [4] using the scVelo velocities with CellRank’s velocity kernel. scVelo only infers a terminate state at the dead-end cells. Note: CellRank offers a method of improving this estimate but incorporating other kernels, but here we just use the velocity kernel to analyze scVelo velocities. **o)** Example latent trajectories of the dynamics inferred by LatentVelo are shown on the UMAP representation of the spliced latent state, showing the divergence of dead-end and reprogrammed cell trajectories, indicating that LatentVelo is correctly inferring the dynamics of reprogramming. 25 dead-end and 25 reprogramming trajectories with 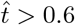 are randomly sampled.

Observing the regulator parameter *z*_*r*_, we see the LatentVelo infers two distinct regimes of dynamics: negative *z*_*r*_ where cells undergo reprogramming, and positive *z*_*r*_ of dead-end cells that do not. This highlights that the regulation parameter *z*_*r*_ can describe different “modes” of the dynamics, and here identifies the differences in reprogrammed vs dead-end cells.

Reprogramming takes a complex path through the data. However in contrast, it becomes clearly apparent in the UMAP plot of the spliced latent space inferred by LatentVelo shown in Figure 3 f)-i). Here, we see the latent space cleanly separates the reprogrammed/dead end cells, generating a latent representation informative of the reprogramming dynamics. This shows that the latent representation of the dynamics inferred by LatentVelo is biologically meaningful. In Supplemental Figure S5, we show in the pancreas, retina, bone marrow, hematopoiesis, intestinal organoid, and hindbrain datasets that in general, the regulatory parameter **z**_*r*_ clearly identifies lineages in this same way, highlighting lineage-dependent dynamics that are not modelled in some other RNA velocity approaches like scVelo.

In Figure 3 j)-m), we show the distribution of *z*_*r*_ for reprogrammed/dead-end cells, and *z*_*r*_ vs latent time, showing a clear separation. We also train a logistic regression classifier on *z*_*r*_ (effectively just scaling *z*_*r*_ to represent a probability), and show a ROC curve, demonstrating that it is predictive of reprogrammed cells. This shows that the parameter *z*_*r*_ inferred by the model corresponds to a meaningful biological feature: the reprogramming outcome.

From just visualizing the embedding overlayed with velocities in 2 dimensions, it cannot be determined if methods infer terminal states corresponding to reprogrammed and dead-end cells. We use CellRank [4] to analyze the estimated velocities of scVelo. We use the velocity kernel, which computes a transition matrix based on the estimated velocities. Using the scVelo velocities, only a single terminal state is obtained in Figure 3 n), corresponding to dead-end cells. This is due to scVelo incorrectly inferring transitions from reprogrammed cells to dead-end cells [4]. CellRank offers a method of improving this estimate by combining the velocity kernel with other kernels, but here we are only interested in the velocity estimates of scVelo. With LatentVelo, we sample 25 trajectories for dead-end and reprogrammed cells in Figure 3 o). We show that these trajectories diverge towards reprogrammed and dead-end states, with no significant transitions from reprogrammed to dead-end cells as there are in scVelo.

This analysis with LatentVelo suggests that the reprogramming fate decision occurs early on in the process, as early as 6 or 9 days (seen at 6 days, but there are very few cells tagged as reprogrammed at 6 days). This was also suggested in the original analysis by Biddy *et al*. [29]. Since LatentVelo is able to infer these early reprogrammed fates without the celltagging information, this suggests that the relevant information for reprogramming is available within the scRNAseq data at early times, and can be extracted with an analysis of the RNA velocity dynamics with LatentVelo.

### C. Latent space dynamics correct for batch effects in RNA velocity and cell states

Batch correction for RNA velocity is currently an unaddressed problem [13]. Since RNA velocity methods operate on gene-space, high-performing batch correction methods [26] that typically output a latent embedding or corrected nearest neighbor graph will not work with typical RNA velocity methods. Another challenge of correcting batch effects in RNA velocity the need to simultaneously correct unspliced and spliced RNA counts, whereas batch correction methods typically operate on a single data matrix.

Since LatentVelo learns dynamics on a latent space, it naturally enables batch correction by learning a shared latent space for the multiple batches. This works in a similarly to existing variational auto-encoder batch correction methods, such as scVI and scANVI [15, 30]. By including batch IDs in the input of the encoder and decoder, and using batch-independent latent dynamics, all batch variation can be handled by the encoder and decoder. This enables LatentVelo to perform batch correction of cell dynamics and cell states.

In Figure 4, we demonstrate LatentVelo’s ability to learn dynamics in the latent space that are independent of batch effects. We use two different simulated datasets from dyngen with strong batch effects. To generate the simulated data with batch effects with dyngen, cell kinetic parameters are sampled from the same distributions in different realizations with the same gene regulatory networks, as suggested in the original dyngen paper [31].

**FIG. 4.**
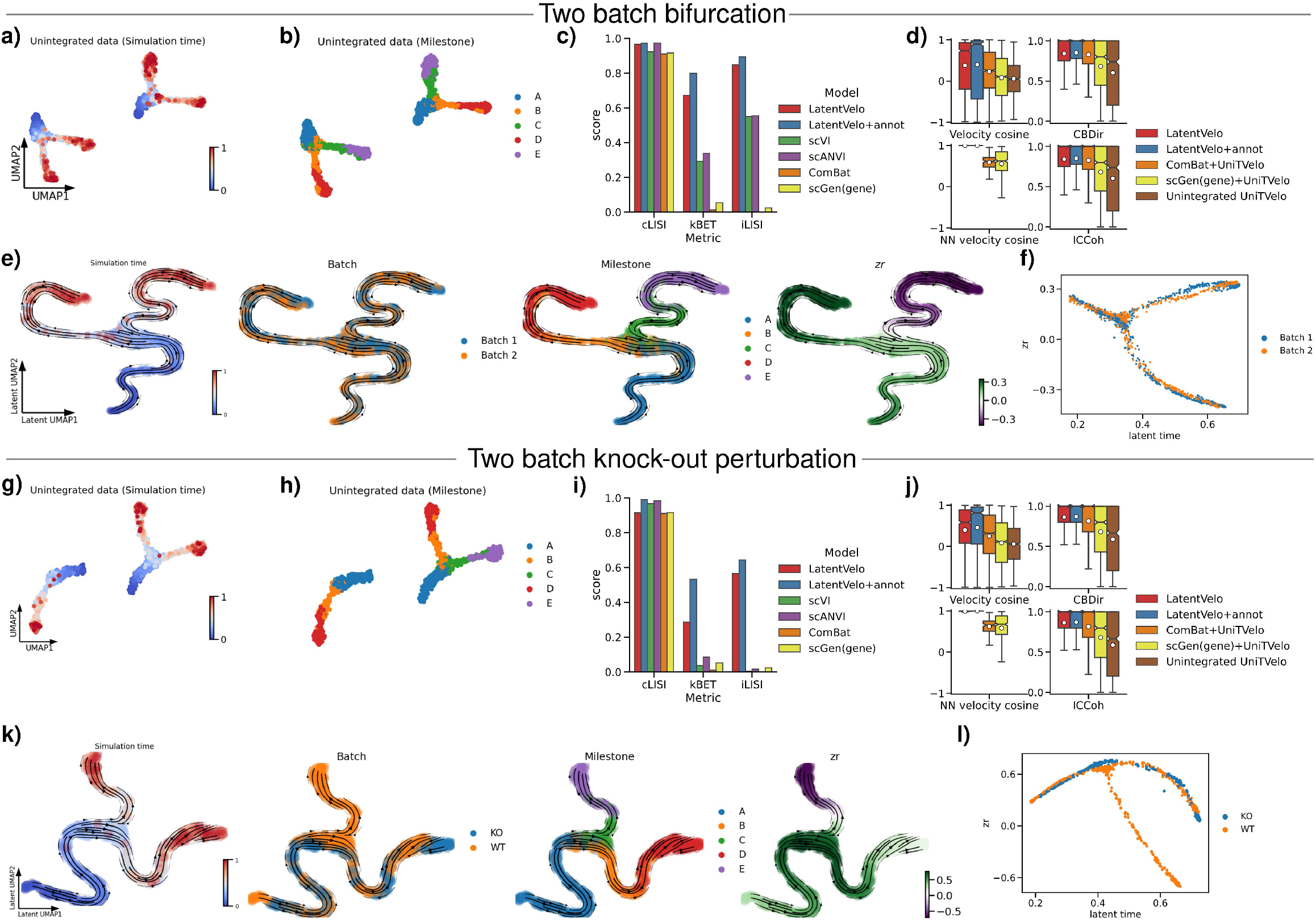
Batch effect correction of dynamics with LatentVelo. **a)** Unintegrated data of a two batch bifurcation, simulated with dyngen. Simulation time shows the direction of differentiation. **b)** Highlighting the “milestone” cells in the data, which define similar clusters of cells between batches and define the direction tested with CBDir scores. **c)** Batch correction (kBET and iLISI) and biological cluster conservation (cLISI) scores for batch correction of cell states (not velocity). Higher scores are better. **d)** Velocity scores for the models. Nearest neighbor (NN) velocity cosine similarity scores measure batch correction of velocity, and the velocity cosine similarity and CBDir scores measure velocity accuracy. Higher scores are better. **e)** UMAP of LatentVelo with celltype annotations (LatentVelo+annot) showing simulation time, batch ID, milestone annotations, and latent regulatory state *z*_*r*_ of spliced latent states at 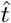. Visually, the data are well integrated. **f)** *z*_*r*_ vs latent time for the two batches, showing batch correction of the regulatory variable on the distinct branches. **g)** Unintegrated data simulated with dyngen. This data contains with two batches with a bifurcation, but for one of the batches a gene-module knockout has been performed, knocking out one of the lineages. (Labelled WT wildtype and KO knockout) **h)** Highlighting the milestone cells, which shows that the C (green) and E (purple) lineage is removed by the knockout. **i)** Batch correction metrics for cell states. Higher scores are better. **j)** Velocity metrics, showing scores for batch correction and accuracy of velocity. **k)** UMAP representation of our LatentVelo+annot spliced latent states. Batches are visually well integrated and follow the correct differentiation trajectory, despite the knockout on one of the batches. **l)** *z*_*r*_ vs latent time for the two batches, showing batch correction of the regulatory variable for the branches. In Supplemental Figure S3, we show UMAP plots for scVI and scANVI on these datasets, highlighting the improvement in batch correction by LatentVelo.

We compare our model with existing batch correction tools for gene-expression (without velocity). Recently a method using traditional batch correction tools that produce a corrected gene count matrix has been suggested for RNA velocity [32, 33]. We compare this method to LatentVelo by using UniTVelo (unified time mode) combined with batch correction tools on gene-space. We have chosen UniTVelo for this comparison because we found it generally performed quite well with default settings on the synthetic datasets, while scVelo (stochastic and dynamical modes) performed poorly even without the included batch effects (Figure 7). We evaluate these methods with metrics of biological cluster conservation (cLISI) and batch correction (kBET and iLISI) [26] for the latent representation of cell-states, as well as with velocity metrics of CBDir and ICCoh and a novel metric of batch corrected velocity (NN velocity cosine similarity), comparing the cosine similarity of velocities for neighboring cells in different batches.

First, we use a simulated dataset with two batches of a bifurcation. In 4a) and b), we show a UMAP representation of the unintegrated data, with the simulation time highlighting the differentiation direction and trajectory milestones indicating similar clusters of cells accross batches for the CBDir metric. In c), we compare our model with traditional batch correction methods (no velocity) for biological cluster conservation (cLISI) and batch correction (kBET, iLISI) of the latent cell states. Our model is the only method scoring highly on all metrics, even when compared to these methods specifically designed for batch correction of gene expression.

In d), we evaluate batch correction of velocity. We compare LatentVelo with UniTVelo used on the unintegrated data, or UniTVelo used with a previously suggested method for batch correction of RNA velocity [32, 33] (see Methods section V H for details). This approach requires the use of any generic batch correction method that output corrected gene-expression matrices, and so we use this method with ComBat [34] (as was originally suggested), and scGen [35] (an auto encoder method). We found that each of these methods fail to properly integrate the batches and have worse performance on velocity metrics than LatentVelo.

In Figure 4e), we show the UMAP plot of the spliced latent state for LatentVelo with celltype annotations (LatentVelo+annot). The model latent space accurately integrates the two batches and RNA velocity accurately follows the direction of differentiation. Both batches are clearly visually integrated in addition to the quantitative metrics. Observing the *z*_*r*_ regulatory parameter at 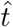, we see that the model accurately describes the lineage dynamics with distinct values of *z*_*r*_ on each lineage, despite the two batches. Additionally, we note that adding celltype annotations to the latent space improves batch correction, compared to other autoencoder batch correction methods that only consider cell states.

In Figure 4g) and h), we demonstrate our approach on two simulated batches of a bifurcation, but now perform a gene-module knockout on one of the batches, eliminating the C (green) and E (purple) milestone cells in this batch. The quantitative metrics in Figure 4i) and j) show that LatentVelo is the only model able to integrate the two batches. In Figure 4k), we show the UMAP representation of the spliced latent states, showing the integration of the batches in the joint latent space. LatentVelo cleanly disentangles the batch effects from the perturbation.

In each of these simulated datasets, LatentVelo is the only model consistently performing strongly for both velocity estimation and batch correction of cell states and velocities. We also see that adding annotated cell-type information to the model is useful to improve batch correction. Additionally, our model performs even better at batch correction of gene expression than the batch correction methods scVI, scANVI, ComBat, and scGen(gene). This is visualized in Supplemental Figure S3. LatentVelo is able to achieve this better performance for batch correction by utilizing the splicing dynamics to not just position similar cells together, but to also align them in time according to the dynamics. All model hyperparameters remain at their defaults for these batch correction tests. We also use default parameters for scVI, scANVI, scGen, ComBat, and UniTVelo.

### D. LatentVelo can infer complex lineage specific gene dynamics

While the main focus of Latentvelo is to infer latent dynamics, we can also estimate the dynamics of single genes. We compute the velocity of single genes in LatentVelo by transforming the latent space velocities into gene-space velocities by utilizing the decoder: 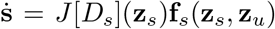, where *J* [*D*_*s*_](**z**_*s*_) is the Jacobian of the spliced decoder evaluated at **z**_*s*_ and **f**_s_ is the latent space velocity. However since the high-dimensional gene space is compressed into a low-dimension latent space, we expect the dynamics of only the best reconstructed genes in the autoencoder to be modelled well.

Previous work has identified Multiple Regime Kinetic genes (MuRK) that show a transcription boost during erythroid development, resulting in a large increase in unspliced RNA [12, 13]. This results in scVelo dynamical and stochastic modes failing to identify the correct velocity direction. This is due to the dynamics of these MuRK genes being poorly modelled by the linear differential equations. We use two datasets containing the development of erythroid cells to demonstrate LatentVelo’s ability to model dynamics with a transcriptional boost. While scVelo fails on these datasets, recent models such as VeloAE, UniTVelo, and VeloVAE have been previously demonstrated to correctly model a transcriptional boost [16, 17, 19].

First, we use a mouse erythroid development dataset [36]. In Figure 5 a)-e), we show that our approach captures the correct differentiation direction from blood progenitors to Erythroid cells, and contrast this with the reversed direction inferred by scVelo.

**FIG. 5.**
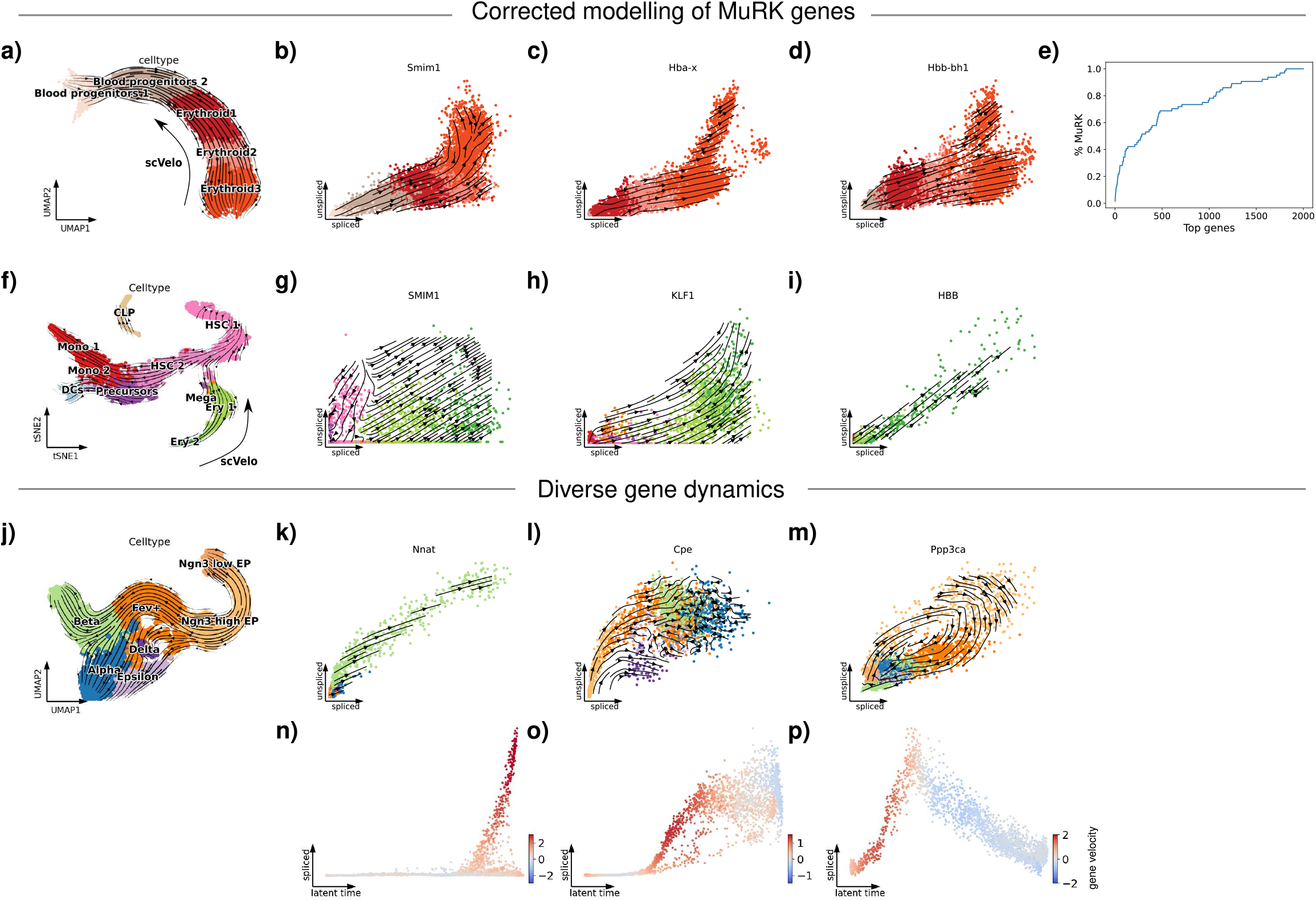
LatentVelo can infer complex gene dynamics. **a)** UMAP plot a mouse erythoid development dataset. LatentVelo velocity arrows on shown on the plot, and the inferred scVelo direction is shown. **b)-d)** We show unspliced vs spliced plots of three genes showing a boost in transcription between Erythroid 2 and Erythroid 3 cells, demonstrating that our model correctly models the velocity of these genes. **e)** We show that many of the best predicted genes in our model are MuRK genes. **f)** t-SNE plot of a human bonemarrow dataset. LatentVelo velocity arrows are shown on the plot, and the inferred scVelo direction for the erythroid lineage is shown. **g)-i)** We show three genes showing a boost in transcription on the erythroid lineage, demonstrating that our model correctly models the velocity of these genes. **j)** UMAP plot of a pancreas endocrinogenesis dataset. **k)-m)** We show three genes showing diverse dynamics for this dataset. **n)-p)** Spliced vs latent time of these three genes with color showing the value of the velocity, showing the sign matches the direction of the dynamics.

Second, we would like to show that even though LatentVelo is modelling a low-dimensional representation of RNA velocity, and not the high-dimensional gene space, we can still infer the dynamics of key critical genes. Unspliced-spliced RNA plots are shown for some of the MuRK genes best reconstructed by the VAE (in terms of *R*^2^ score), b) Smim1, c) Hba-x, and d) Hbb-bh2. These MuRK genes show a transcriptional boost during the transition between Erythroid 2 and Erythroid 3. In comparison to scVelo where a reversed velocity direction is seen [12], our approach correctly models the velocity direction for these genes.

MuRK genes previously identified in this dataset contain archetypal red blood cell genes essential for red blood cell function [12]. We demonstrate that LatentVelo utilizes these critically relevant genes by showing the percent of top reconstructed genes in terms of *R*^2^ score that are MuRK genes in Figure 5 e). This shows that even though our model latent space is compressing the 2000 highly variable genes to 20 dimensions, it is still capturing these critical genes.

Note that since this dataset includes 3 batches, we include the batch ID in the encoder and decoder for batch correction with LatentVelo. This is most clearly seen in the Hbb-bh1 gene unspliced vs spliced plot (Figure 5 d), where two separate blobs of Erythroid 3 cells (dark orange) are seen due to belonging to separate batches. Due to our model including batch correction, when plotting these raw unintegrated counts we see the model accurately predicts different velocities on each of the batches.

We also test our method on a human bonemarrow dataset [37]. Similar to the mouse erythroid dataset, scVelo has many problems on this dataset, in particular showing incorrect differentiation direction on the erythroid lineage and showing differentiation towards hematopoetic stem cells (HSC). In Figure 5 f) we show that our method accurately models the direction of differentiation, from HSC to the erythroid lineage, and to the monocyte (Mono 1,2) and dendridic (DCs) lineages. We also highlight some example genes, where our model captures the transcriptional boost behaviour of the g) SMIM1, h) KLF1, and l) HBB genes on the erythroid lineage. Additionally, in the SMIM1 gene plot, we show that LatentVelo infers distinct behavior for this gene on the different lineages: a transcriptional boost of the gene on the erythroid lineage and a repression of the gene on the other lineages.

The more general case of time-dependent rates is explored in Supplemental Figure S6. We show that our model is robust to increasing/decreasing transcription, splicing, and degradation rates vs time with simulations.

In Figure 5 j)-m) we show a 3 different genes with a diverse spectrum of dynamics. In k), the Nnat gene is only expressed in the Beta lineage, in l) the Cpe gene is expressed in all lineages at different amounts, and in m) the full cycle of induction and repression of the gene Ppp3ca is seen. In Figure 5 n)-p) we show that the value of the velocity for these genes correctly corresponds to the direction of the dynamics.

### E. LatentVelo infers cell fate trajectories in large multi-lineage systems

In the pancreas dataset in Figure 2, fibroblast reprogramming dataset in Figure 3, and Supplemental Figure S5, we demonstrated that LatentVelo can learn dynamics in datasets with multiple lineages. However, these datasets had at most 3-4 lineages. We now test LatentVelo on a large-scale many-lineage system.

In Figure 6 we show a dataset of mouse gastrulation, showing development from pluripotent epiblast cells into the ectodermal, mesodermal, and endodermal progenitors of major organs [36]. Note, this is the full dataset from where the erythroid cells from Figure 5 come from (seen in the upper right of the UMAP plot in Figure 6a with the same colors). On this large scale dataset, we have increased the dimension of the latent **z**_*s*_ and **z**_*u*_ states to 70 and used a 6-dimensional **z**_r_.

**FIG. 6.**
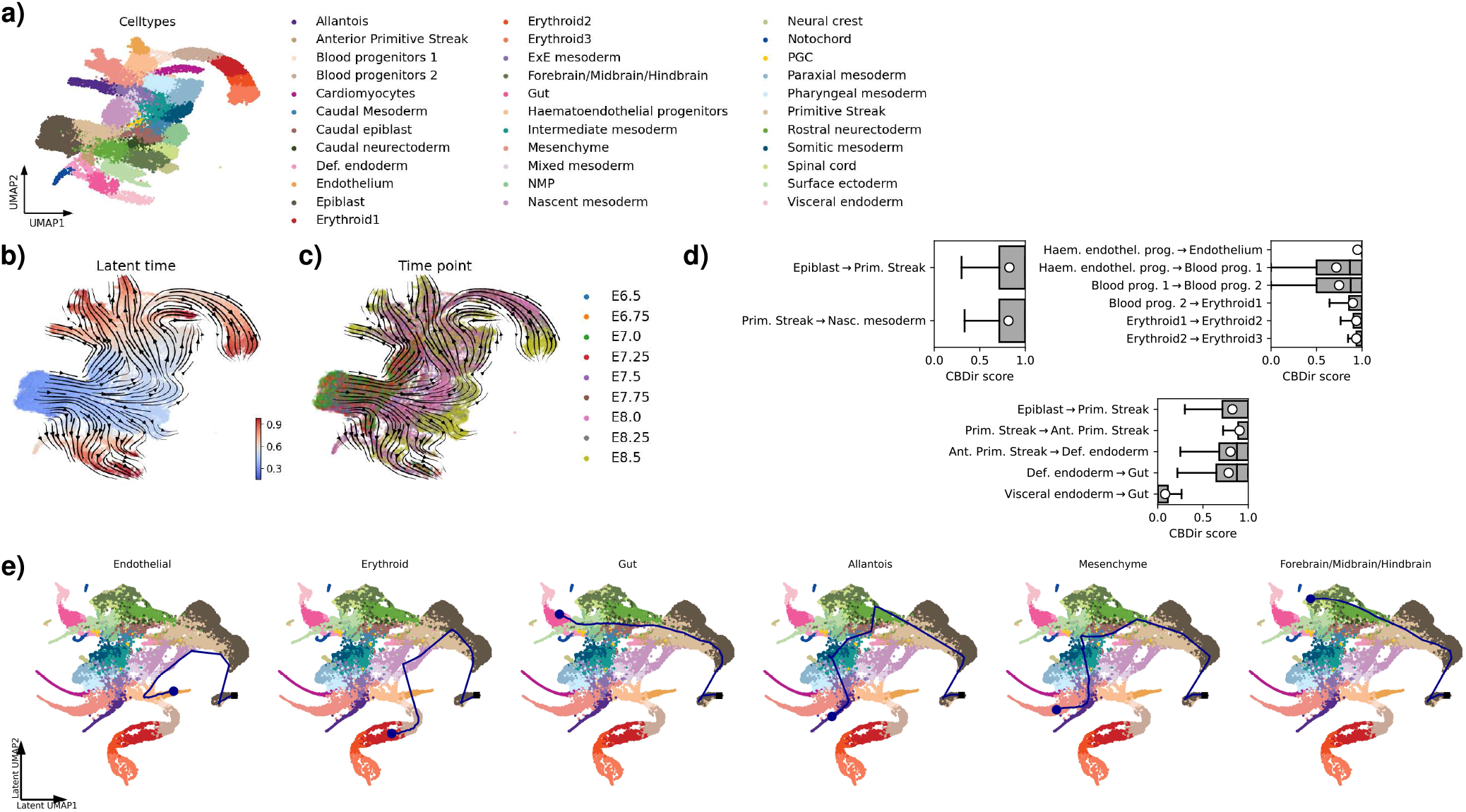
Inferring multiple-lineages in mouse gastrulation. **a)** UMAP plot showing celltypes during mouse gastrulation. **b)** LatentVelo latent time and **c)** experimental time points with inferred Latentvelo velocity. **d)** Some CBDir celltype transition scores estimated on the inferred latent space, highlighting the inferred transitions. **e)** Inferred latent trajectories for examples of endothelial, erythroid, gut, allantois, mesenchyme, and forebrain/midbrain/hindbrain cells.

In Figure 6 c) putative developmental directions can be seen by observing the later time point cells (E8.5), and our model’s velocity and latent time estimates generally agree with these time points in Figure 6 b). We also highlight some CBDir transition scores between celltypes in d), showing the model infers transitions from epiblast to mesoderm, the development of epithelial and erythroid cells, and the development of the gut from endoderm cells. One particular transition that we don’t see is the convergence of visceral endoderm cells and definitive endoderm cells to gut cells that was originally identified [36], we only see the transition from definitive endoderm cells to gut cells. We suspect this is due to LatentVelo assuming a single initial state **z**_0_.

In Figure 6 e), we show inferred trajectories for example endothelial, erythroid, gut, allantois, mesenchyme, and forebrain/midbrain/hindbrain cells. These trajectories give a more clear visualization of the dynamics as compared to the velocity field on the gene-space UMAP plot, where disentangling the many occurring transitions is difficult. Inferred trajectories start as epiblast cells and transition to the primitive streak, then branch to the variety of celltypes.

This dataset demonstrates LatentVelo’s ability to learn dynamics on large multi-lineage systems with many different cell types.

### F. Quantitative benchmarking on synthetic and real datasets

We primarily compare our model with scVelo dynamical and stochastic modes [11]. Note: scVelo stochastic mode is an updated version of the Velocyto steady-state model, where regression is done for the first 2 moments of the dynamics instead of just the mean [11]. Other recent RNA velocity methods have been developed, such as UniTVelo [19], DeepVelo [18], VeloVAE [19], and VeloVI [38]. We do not compare with these methods due to their recency and the variety of hyperparameters needed to be adjusted to fairly compare the methods.

We use dyngen [31] to generate synthetic datasets of 5000 cells each with a variety of developmental structures: linear (51 genes), bifurcation (65 genes), trifurcation (81 genes), and a binary tree (89 genes). In Figure 7 a)-c) we show results on these synthetic datasets. We show the gene space velocity cosine similarity and 50 principle component (PC) cross-boundary directedness score (CBDir) and inner cluster coherence score (ICCoh). The boxes show the interquartile range over the aggregated cells of all celltypes, the notch shows the median, the white point shows the mean, and the whiskers show 1.5× the interquartile range. Since the exact velocities in the simulation are noisy, we average over 100 nearest neighbors when computing the gene space cosine similarity between estimated and exact velocities. Otherwise, the whiskers for velocity cosine similarity extend the entire plot due to noise in the simulation. Separately plotting these “incorrectly” predicted cells shows their simulated velocity going opposite the direction of differentiation. For this reason, we compare with local averages of velocity.

**FIG. 7.**
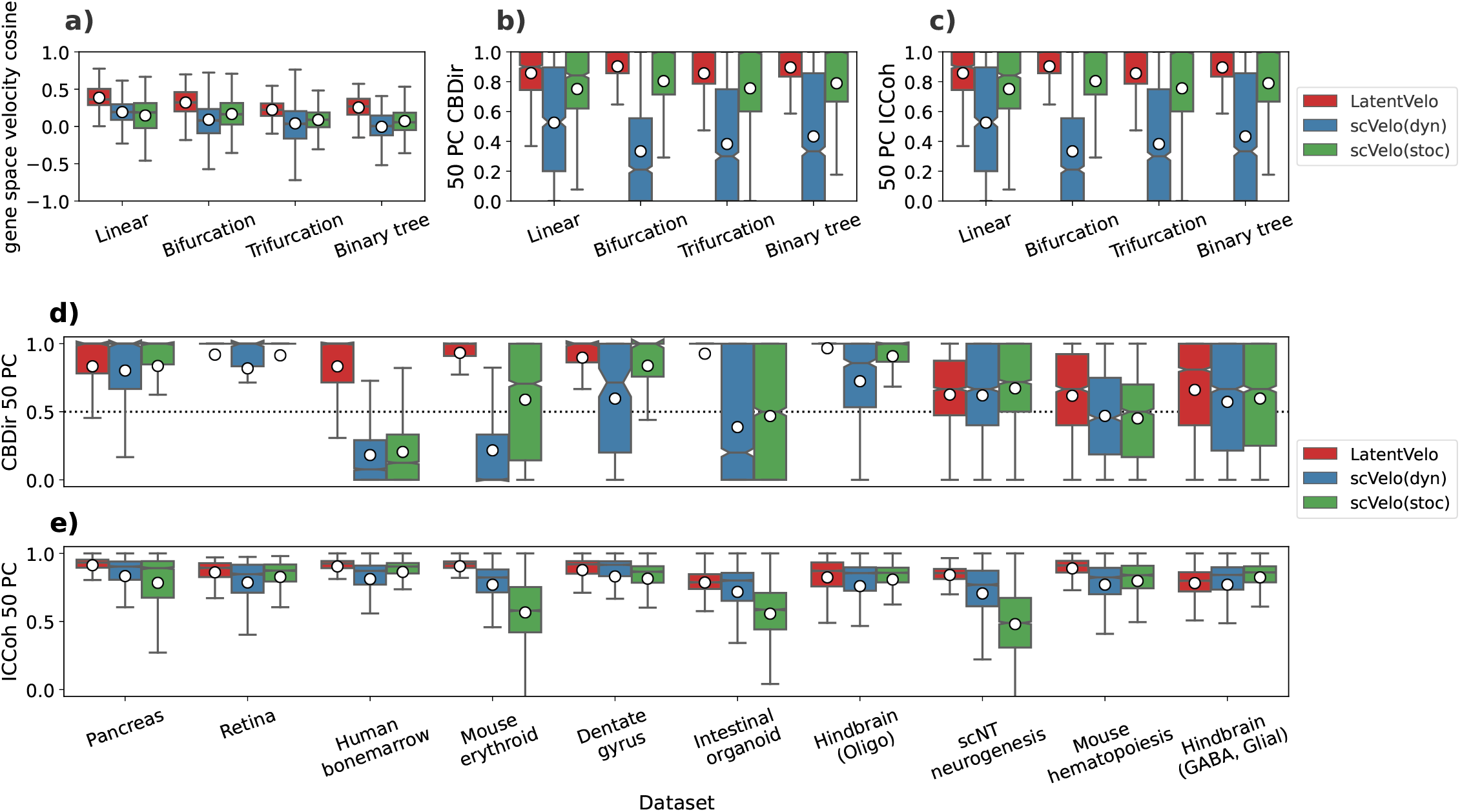
Quantitative benchmarking on synthetic and real data. **a)-c)** Quantitative metrics for the synthetic datasets. We show the a) cosine similarity between estimated and exact simulated velocity on the full gene space, **b)** Cross-Boundary Directedness score (CBDir) on the space of 50 principle components (PC), and **c)** Inner-Cluster Coherence score (ICCoh) on the space of 50 PC. **d)-e)** Quantitative metrics for real datasets. We show **d)** CBDir and **e)** ICCoh scores on the space of 50 PC. Higher scores for all metrics are better.

With synthetic data, we show LatentVelo performs much better than scVelo dynamical and stochastic modes in Figure 7. For LatentVelo and all of the comparison models, hyperparameters and settings are kept at their defaults on these synthetic datasets. We also test our model on increasing/decreasing transcription, splicing, and degradation rates vs time with simulations from the scVelo linear differential equations in Supplemental Figure S6.

In Figure 7 d)-e), we evaluate on 10 real datasets with known trajectory directions, and compare with scVelo stochastic and dynamical modes. Our model consistently scores well for the CBDir and ICCoh scores, above scVelo. One limitation to the CBDir metric is that noisy boundaries between annotated celltypes can obscure the evaluation of velocity directions, and artificially lower scores. In particular this is seen in the scNT, mouse hematopoiesis, and hindbrain (GABA, Glial) datasets, where velocity follows the expected directions but noisy celltype boundaries lower the CBDir scores.

We also evaluate LatentVelo’s ability to model the dynamics of separate lineages with the regulatory parameter **z**_*r*_. We train a simple logistic regression classifier on **z**_*r*_ to predict separate lineages, obtaining classification accuracy between 0.9 and 1 for all datasets (shown in Supplemental Figure S5). This shows that the lineages are clearly separated and modelled with distinct dynamics.

Note that while we have used 50 principle components to evaluate cell-type transitions because we can do a common benchmark between all methods, a natural embedding to use when evaluating cell-type transitions with LatentVelo is just the latent embedding inferred by the model.

For these synthetic and real datasets we keep all model hyperparameters at their default values, except the dimension of **z**_*r*_ and the dimension of **h**. The dimension of **z**_*r*_ is by default set based on the number of expected lineages, and we set *z*_*r*_ to be the number of expected lineages minus one, except if we only expect 1 lineage where we use 1 dimension. We have found this to be a good heuristic. The dimension of h is set to be the same as **z**_*r*_, unless there is poor agreement between 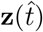 and 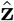, then it is increased (alternatively the dimension of the latent state can be increased, see Supplemental Figure S4). Regularization strength is set to the default value 0.1 for all synthetic datasets and real datasets. For the scNT-seq dataset, the data is very noisy, so we restrict the model to only a small subset of velocity genes (see Methods section V B). However, more work needs to be done exploring these settings of LatentVelo, as well as the settings of other models.

Since all of the parameters of the neural networks in LatentVelo need to be initialized and training is done in random mini-batches, there is some stochasticity in training. For most datasets the model is robust to this, however is some cases LatentVelo infers incorrect reversed velocities on a fraction of the random seeds. A similar problem was encountered with UniTVelo, and was addressed by initializing the latent time with a diffusion pseudotime based on a specified known root celltype, or fitting the model multiple times and selecting the one that is consistent with any prior knowledge [17]. In LatentVelo we mitigate this problem on the mouse gastrulation, fibroblast reprogramming, and hindbrain GABA datasets by including a correlation in the loss function between inferred latent time and experimental timepoints for the first few epochs of training (see Methods section V D). We have found that reversed velocities result from the initialization of the model, and this can be corrected early by simply influencing the correct direction with experimental time-points. This also is a problem on the scNT, intestinal organoid, and bonemarrow datasets, where experimental timepoints are not available. In these cases, the correct direction must be chosen out of multiple random seeds, or the “root” cells need to be specified (see Methods section V D).

We believe this issue with randomly fully reversed velocity fields occurs due to a lack of genes showing cycling induction to repression behaviour. The result is monotonic gene dynamics, where the direction is ambiguous. These problems occur on datasets where scVelo also has problems. In LatentVelo offer the ability to specify root cells to correct for this issue, or use experimental time points.

## IV. DISCUSSION

We have introduced LatentVelo, a new model of latent cell developmental dynamics. We have demonstrated that our model is accurate and robust on a wide variety of synthetic and real datasets, outperforming the currently most widely used approach, scVelo [11]. LatentVelo is the first method to batch correct RNA velocity by inferring velocity on a latent embedding, and additionally performs better at batch correction of gene expression than other methods that only attempt to batch correct gene expression and ignore dynamics. In addition to just cell velocities, LatentVelo infers the trajectory a cell took to its current state, which is useful for improving the interpretation of dynamics. LatentVelo can infer distinct RNA velocity dynamics on separate lineages by learning a latent regulatory parameter. We also show that the latent embedding and the latent regulatory parameter can represent biologically meaningful features.

Previous RNA velocity methods have inferred velocity on gene-space, with the only exception being VeloAE, which only modelled steady-state dynamics [16]. LatentVelo embeds gene expression into a latent space, with the time-progression of gene expression described by latent dynamics. Since the latent embedding and latent dynamics are trained together, this allows us to infer dynamics-informed embeddings of gene expression, which is a dimensional reduction of cell states, informed by the dynamics of the system. Shown with the fibroblast reprogramming dataset [29], the UMAP representation of this latent space allows clear visualization of the separate trajectories taken by reprogrammed or dead-end cells. This opens up a new method of studying and characterizing the dynamics of cell differentiation, through the features represented in this latent space.

LatentVelo’s main application is describing complex developmental dynamics and inferring trajectories in a low-dimensional latent space. Despite modelling dynamics in this low-dimensional latent space, LatentVelo can still model the dynamics of single genes by transforming velocities back to gene-space with the decoder. In Figure 5 we showed that LatentVelo can model the dynamics of MuRK genes in erythroid development and a diverse set of gene dynamics in pancreatic endocrinogenesis. However since the high-dimension gene space information is highly compressed, this can only be reliably done for the genes that are well reconstructed by the autoencoder. This is one limiting feature of LatentVelo in comparison to other approaches that directly model dynamics on gene space.

Interestingly, we have found scVelo stochastic mode, which computes RNA velocity based on a linear regression between unspliced and spliced RNA at an assumed stead-state, performed much better than scVelo dynamical mode. We believe this highlights the issues with the simple linear differential equation with constant rates approach of scVelo dynamical mode. The recent methods DeepVelo [18], UniTVelo [17], VeloVAE [19], and our approach LatentVelo all have different approaches to addressing this issue by generalizing the simple linear dynamics.

Since LatentVelo is a variational auto-encoder, we can sample the latent space to generate uncertainty estimates of latent times and velocities. This is shown in Supplemental Figure S7, where we show uncertainty is largest near branching between multiple cell-types. This method of uncertainty estimation is distinct from the method of scVelo [11], which used the consistency of the velocity estimates of neighboring cells. Recent work has extensively explored these types of uncertainty estimates from variational Bayesian models of RNA velocity [38, 39].

LatentVelo can be easily extended to multi-omics. In the Supplemental Information, we show an example of this with combined scRNA-seq and ATAC-seq. This extension is done by adding a new variable to the structured latent dynamics **z**_*c*_, corresponding to the latent representation of chromatin accessibility, and modelling the regulation of chromatin by **z**_*r*_ and the regulation of transcription by **z**_*c*_ and **z**_*r*_. This demonstrates a key feature of LatentVelo: the ability to build *general* structured latent dynamics. This is presented as an example, we leave the more detailed analysis of multi-omic systems and exploration of other forms of structure in the dynamics to future work.

LatentVelo addresses many of the challenges raised in a recent review paper of RNA velocity [13]: multi-modal omics (see Supplemental Information), multi-variate dynamics, batch correction, lineage/time dependent rates, and implicit gene selection by embedding in the latent space. Two challenges not addressed are (1) stochastic dynamics, and (2) normalization. We see potential ways to address these challenges with simple modifications to our model: (1) for stochastic dynamics we can replace the latent ODEs in our model with SDEs [40], and (2) we can approach normalization in a similar way as scVI [30]; including normalization factors as latent variables to infer. Modelling SDEs instead of ODEs is a clear next direction for LatentVelo, in particular by using a Bayesian approach to inferring the latent SDE [40], we can simultaneously learn the prior SDE describing the full population of cells as well as the conditional SDE describing the dynamics of each cell conditioned on a particular branch (as we have done with the parameter h). This would allow the use of the model for perturbed inputs – estimating future developmental trajectories for individual cells, rather than just the past trajectories of cells as we have done here. Existing SDE models do not learn the latent embedding as LatentVelo does, nor do they incorporate splicing dynamics [41].

There are limitations to the evaluation of RNA velocity on real datasets. We have used a modified form of the CBDir metric [16, 17], which quantifies the probability of each cell transitioning to the expected celltype. A substantial limitation of this approach is that it relies on accurate discrete annotations of cell-types, which can cause issues on the boundaries between celltypes. Additionally, the CBDir metric is poor at identifying the absence of a transition between two cell types because it does not look at the strength of transitions, only the direction of velocity. Further work needs to be done on the development of quantitative metrics for RNA velocity.

A limitation of LatentVelo is that by default we assume a single initial latent state from which the trajectory of all cells start from, and that each observed cell can be reached by a continuous trajectory from this initial state. For example in the full mouse gastrulation dataset (Figure 6), we do not see the expected convergence of visceral endoderm cells and definitive endoderm cells to gut cells [36]. Since each cell needs to be reached by a continuous trajectory, we may encounter issues in situations with sparse cell clusters with few observed transient cells connecting trajectories. We believe that this occurs in the variety of different sparse cell clusters in the dentate gyrus dataset [11]. In these cases the model still infers generally accurate velocity directions for a given cell (as seen in the benchmarking in Figure 7), but latent trajectories do not start from expected initial states. For example, on the dentate gyrus dataset, we see the expected transitions on the granule lineage, radial glia to astrocyte transitions, and oligodendrocyte progenitors to oligodendrocyte transitions, but LatentVelo estimates the disconnected microglia cluster as having the lowest latent times, so inferred trajectories all incorrectly start from this cluster.

While we have shown LatentVelo is accurate on many datasets, for some it requires additional hyperparameter adjustment by increasing the dimension of the latent states, using the celltype annotated model, or restricting to only a subset of genes. Indeed, all recent improvements to RNA velocity also have multiple settings and hyperparameters, in addition to preprocessing steps [14]. While we have discussed some heuristics to choosing the hyperparameters of LatentVelo, more work needs to be done in this area.

## V. METHODS

### A. LatentVelo

We take spliced and unspliced counts per cell as input and embed into a latent space 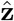 with an encoder neural network. A separate encoder is used for spliced and unspliced counts, partitioning the latent space as 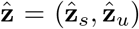. Our model is trained as a Variational Autoencoder (VAE), where we use a standard normal prior on the latent space 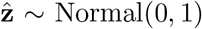. We also use an encoder to estimate a latent developmental time with a logit-normal prior *t* ∼ LogitNormal(0, 1).

Dynamics on the latent space are described by neural ordinary differential equations,

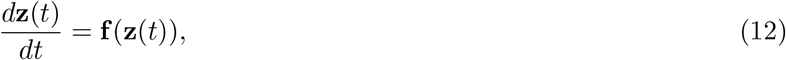

where **f** is a neural network describing the velocity field of the dynamics. The spliced component of this velocity field represents latent RNA velocity. The structure of **f** is decomposed into the 3 components described in Figure 1, **f** = (**f**_*u*_, **f**_*s*_, **f**_*r*_).

These dynamics are coupled to the auto-encoder by matching the ODE solution 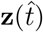 at time 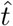 with the encoded latent state 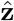.

We use an approximate posterior factorized as 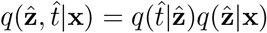, where **x** = (**s, u**) are the observed spliced and unspliced counts. Separate encoders for spliced and unspliced latent states are used, such that 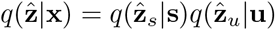. Additionally, we also use separate decoders, *p*(**s, u**|**z**) = *p*(**s**|**z**)*p*(**u**|**z**). Since the data are very noisy, we find that we cannot use a count-based distribution such as a negative-binomial, so instead use a Gaussian with mean functions *μ*(**z**) = (*μ*(**z**_*s*_), *μ*(**z**_*u*_)) with smoothed and normalized counts following the scVelo preprocessing procedure [11]. Our model is trained with the loss:

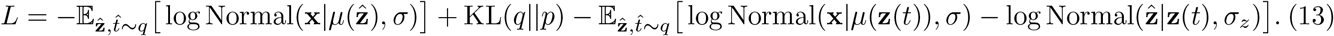

Expectations are computed by sampling 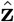 and 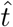 from the encoder. In Methods sections V B and V D we discuss further terms in this loss function regularizing the trajectory direction.

The first two terms of this loss are the standard negative evidence lower bound of a VAE [23, 24], representing the expectation of the negative log-likelihood with a Gaussian, and the KL divergence between the posterior 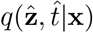 and the prior, Normal(0, 1) × LogitNormal(0, 1). The two remaining terms include a loss for the decoded solution of the dynamics 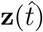, and a term penalizing the distance between the encoded latent state 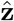 and the latent state estimated by the dynamics 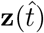.

By default, we use a 20-dimensional latent state for each observed component (spliced, unspliced) and use ELU activations throughout. We follow VeloAE [16] and use a encoder structured as an initial fully connected dense neural network with 1 hidden layer (of size 25 by default) as the first part, and then use two graph convolution layers using a 30 nearest-neighbor graph computed on the 50 principle components (same as was used for smoothing) as the similarity graph for the second part. This enables the encoder to use information about nearest neighbors when computing the latent embedding. In the case of batch correction, we do not use the graph convolution layers and instead just use a fully connected dense neural network with 2 hidden layers.

The encoders for **h** and 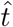 are also graph convolutional networks with 2 layers of graph convolutions. The differential equation derivative functions **f**_*u*_ and **f**_*r*_ have a single hidden layer of size 25. **f**_*s*_ is linear in **z**_*s*_ and **z**_*u*_. By default we also use a linear decoder, and use separate linear decoders per batch for batch correction (with the option for a fully connected neural network). The dimension of **z**_*r*_ is chosen based on the expected number of branches. For one or two expected branches, we use 1 dimensions, for 3 expected branches we use 2 dimensions. In general, we take the dimension of **z**_*r*_ to be 1 less than the number of expected branches.

We use 90% of the data for training, and use the other 10% as a validation set to monitor training progress. We train for 50 epochs, and use the model at the epoch with lowest mean-squared error on the validation set. By default we train with the Adam optimizer with a learning rate of 0.01 and a batch size of 100. In large datasets (e.g. mouse gastrulation or Fibroblast reprogramming), we increase the batch size to 1000. In scenarios with exploding gradients resulting in failed training, we use gradient clipping to stabilize training. We increase the KL divergence weight in the VAE from 0 to 1 linearly over the first 25 epochs.

The model is implemented in *pytorch* utilizing the *torchdiffeq* package for neural ODEs [25, 42], which critically allows computing gradients of the inferred latent time-points 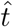 in the solver, rather than marginalizing over 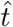 by integrating. We found the marginalization approach to be challenging.

### B. Enforcing splicing direction

The linear ODEs describing transcription, splicing, and degradation are,

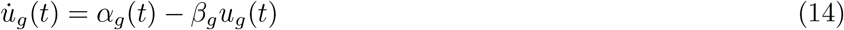

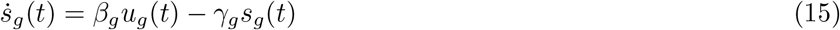

per gene *g*. These equations enforce the casual relationship of splicing between unspliced and spliced RNA. Generalizing, this same effect can be achieved with 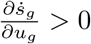 and 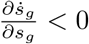.

To constrain the direction of splicing to accurately represent the biology (unspliced to spliced), we need to enforce 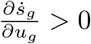 and 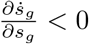 for each gene *g*. However, enforcing this constraint for general forms of the velocity 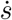 is difficult, since we need to compute the Jacobian, which is a matrix of size 2*N* ^2^, where *N* is the number of genes. Computing this would require back-propagation through the entire model back to the input, which is computationally infeasible. Instead, we can use a much faster approach to enforce this direction by using the correlation between velocity and spliced and unspliced counts.

We weakly regularize by the correlation between gene-space velocity and input data (similar to DeepVelo [18]),

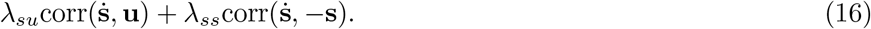

By default we take *λ*_*su*_ = *λ*_*su*_ = 0.1. The goal with this term is to just weakly regularize the direction, rather than match the strict linear dependence seen in other models. To compute gene-space velocity we compute the time-derivative of the transformation with the decoder, 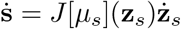, where *J* [*μ*_*s*_] is the Jacobian of the spliced decoder. Similarly we can compute the unspliced gene-space velocity, 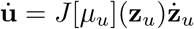. Note, the Jacobian here is taken with respect to the latent space which has a smaller dimension, and this is a Jacobian vector product, which are easily computable by backpropagation in comparison to the Jacobian discussed above. This regularization can be done per celltype, only including the same type cells in the correlation.

We only apply this regularization to genes that show significant splicing dynamics. These “velocity genes” are identified in a similar way to UniTVelo and scVelo [17]. We fit a linear regression between spliced and unspliced data per gene, then only select genes with ***R***^2^ score above 0.05 and below 0.95. In cases of genes with very high ***R***^2^, all cells lie on a straight line in the *u* vs *s* plane, showing no splicing dynamics. In cases of genes with very low ***R***^2^, cells are scattered uniformly in the *u* vs *s* plane, showing no splicing dynamics. Genes with an extreme ratio of standard deviations between unspliced and spliced were also filtered so that 0.3 ≤ *σ*_*u*_*/σ*_*u*_ ≤ 3, given that this may be the result of error in reading unspliced counts.

### C. Incorporating cell-type annotations

We follow scANVI [15], and incorporate cell-type annotations by modifying the prior. We introduce the new latent state ŵ, and use the new approximate posterior,

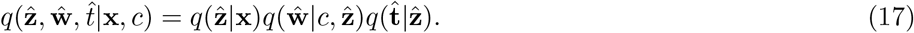

We place a standard normal prior on w, and model 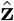 and 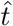 as functions of w and the cell-type annotations *c* instead of placing a standard normal/logitnormal priors; 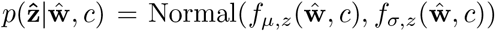 and 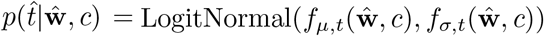.

The effect of this modification is to allow the priors on 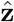 and 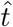 to vary with cell-type, preventing over-regularizing and eliminating biologically relevant cell-type clusters for the sake of matching the restrictive prior.

### D. Incorporating experimental time points or root cells

We can include a regulation term to enforce a positive correlation between inferred latent time and experimental time when available, 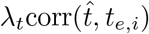, where *t*_*e,i*_ is the experimental time point for the *i*th cell. The weight of this correlation can be linearly decayed over the first few epochs to fix reversed velocities.

In scenarios where no experimental time-points are known, a “root” celltype can be used in reversed velocity scenarios. In these cases we can regularize the estimated latent time 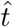 to be minimal for the specified root celltype.

### E. RNA velocity and latent time uncertainty

To estimate uncertainty in latent time, we sample from the inferred posterior distribution 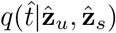 and compute the standard error of the mean.

To estimate uncertainty in latent RNA velocity, we sample from the inferred posterior distribution of states 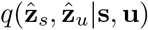 and compute latent RNA velocity for each sample 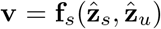. We then compute the average cosine similarity of all pairs from these samples as the consistency of velocity,

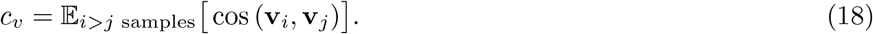

### F. RNA velocity metrics

When ground truth velocities are known (synthetic data), we compare estimated velocities by computing the cosine similarity on gene space.

When ground truth velocities are not known, we use known cell-type transitions with the Cross-Boundary Directedness metric [16, 17].

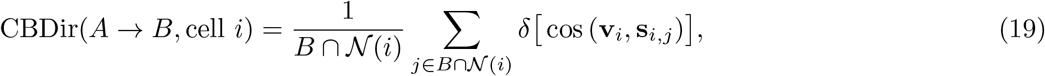

where **s**_*i,j*_ = (**s**_*i*_ − **s**_*j*_)*/*sign(**s**_*i*_ − **s**_*j*_) and 𝒩 (*i*) is the neighborhood of cell *i*. Since the boundaries between cells can be noisy in high-dimensional gene space, we use 50 principle components when computing this score.

We also use the In-Cluster Coherence (ICCoh) metric [16, 17] to evaluate the coherence of velocities within a cluster or cell-type. This score is computed per cell by the average cosine similarity of neighboring cells in the same cluster/celltype,

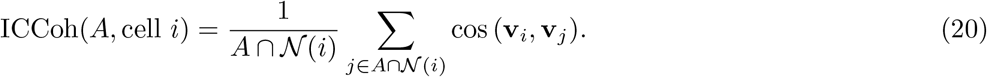

Similarly, we also use 50 principle components for this score.

### G. Batch correction metrics

To evaluate batch correction we use the kBET and iLISI metrics [26]. We also use the cLISI metric to evaluate biological cluster conservation [26], ensuring that cell-type clusters are not erased by the batch correction. These metrics are computed with the *scib* package [26].

kBET evaluates the batch composition of the nearest neighbors of a cell, which should match the overall batch composition for a particular cell-type for good batch correction. iLISI and cLISI measure the nearest-neighbor graph structure, evaluating batch mixing (iLISI) or cell-type separation (cLISI).

To evaluate batch correction of RNA velocity, we measure the cosine similarity of nearest neighbor cells in different batches.

### H. ComBat and scGen RNA velocity batch effect correction

We follow the approach taken by Hansen and Ranek *et al*. [32, 33] to compare our approach for batch effect correction of RNA velocity. Since traditional RNA velocity methods require cells in gene-space, only batch correction methods that return a corrected gene matrix can be used. Here we use ComBat [34] and scGen (using the corrected gene output) [35]. ComBat is run from the *scanpy* package [43].

Since we need to simultaneously correct spliced and unspliced counts, batch correction is performed on the sum of these counts. We define the sum matrix *M* and the ratio matrix *R*,

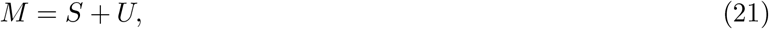

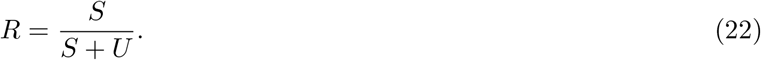

Batch correction is performed on M. For ComBat, we first log(1 + *x*) transform this matrix. For scGen, we normalize then log(1 + *x*) transform.

After batch correcting *M* to get the corrected matrix 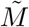, we invert these transforms and then multiply *R* or 1 – *R* to recover corrected spliced and unspliced matrices.

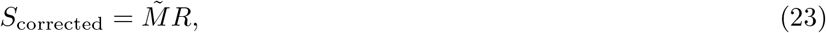

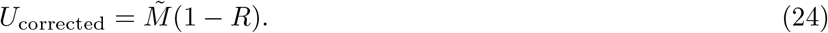

Then RNA velocity is estimated as before with the same pre-processing steps.

scGen is run with default settings, except we set the dimension of the latent state to be the same as our model, 20.

### I. scVI and scANVI batch correction

scVI and scANVI are run from the scvi-tools package [44]. We run scVI with a negative binomial gene-likelihood, and the same latent dimension as our model, 20. scANVI is trained by starting with the pre-trained scVI model and training for an additional 100 epochs.

### J. Comparison velocity models

scVelo stochastic and dynamical modes are run with default settings. UniTVelo is run with the defaults IROOT = *None* and R2 ADJUST = *True*.

When we compute the transition matrix to embed RNA velocity in 50 dimensional PC space or 2 dimensional UMAP or tSNE for plotting, we use scVelo’s function scvelo.tl.velocity graph with default settings.

### K. Datasets

For each dataset we select genes with at least 20 cells with non-zero unspliced and spliced counts, and from these genes select the top 2000 highly variable genes using scVelo preprocessing. We apply the transformation *log*(1 + *x*) before computing principle components and smooth data by averaging over the 30 nearest neighbors in 30 dimensional principle component space with the scVelo function scvelo.pp.moments. Input variables to the model are then scaled to standard deviation 1. For each dataset, we use the provided celltype annotations used in the original publication.

#### Dyngen synthetic datasets

We generate synthetic datasets with dyngen [31]. For each dataset in Figure 7, we simulate 5000 cells. We set 15 target genes and 15 housekeeping genes, and use the number of transcription factors required for the trajectory backbone. We use the backbones “linear”, “bifurcating”, “binary tree” with 2 modifications, and “trifurcating”. We set *τ* = 0.01, census_interval=1, and use 100 simulations. For the CBDir metric, we use the milestones defined by dyngen as transitions. We set the dimension of **z**_*r*_ to be 1 on the linear and bifurcation datasets, and 2 on the trifurcation and binary tree datasets. We set the dimension of **h** to be 1 on the linear and bifurcation datasets, and 3 on the trifurcation and binary tree datasets.

#### scVelo linear model synthetic datasets

We use the scVelo linear differential equations to simulate datasets with time-varying alpha (transcription rate), beta (splicing rate), gamma (degradation rate). We simulate 500 cells with 30 genes. By default, we set *α* = 5, *β* = 0.5, *γ* = 0.5. For time-varying rates, we select 5 genes to be time-dependent. We set the dimension of **z**_*r*_ to be 1 on these datasets.

#### Pancreatic endocrinogenesis

Mouse pancreatic cells sampled at E15.5 [27]. In Figure 7 Initially cycling population is removed to focus on differentiation into terminal cell types. This dataset is downloaded from the CellRank package [4]. The transitions tested with CBDir are (Ngn3 low EP → Ngn3 high EP), (Ngn3 high EP → Fev+), (Fev+ → Delta), (Fev+ → Beta), (Fev+ → Epsilon), (Fev+ → Alpha). We set the dimensions of **z**_*r*_ and **h** to be 3 and on this dataset, and use a latent dimension of size 30. We use the celltype annotated model for this dataset.

#### Mouse hematopoiesis

Data is from [45], and processed data is downloaded from https://zenodo.org/record/6110279 [32]. The transitions tested with CBDir are (LTHSC → MPP), (MPP → LMPP), (MPP → CMP), (CMP → GMP), (CMP → MEP). We set the dimension of **z**_*r*_ and **h** to be 1 and 2 on this dataset.

#### Mouse retina development

Data from the Kharchenko lab http://pklab.med.harvard.edu/peterk/review2020/examples/retina/. The transitions tested with CBDir are (Neuroblast → PR), (Neuroblast → AC/HC), (Neuroblast → RGC). We set the dimension of **z**_*r*_ and **h** to be 2 and 2 on this dataset.

#### Dentate Gyrus development

Mouse Dentate Gyrus development at two time points P12 and P35 downloaded from the scVelo package [11]. The transitions tested with CBDir are (OPC → OL), (Radial Glia-like → Astrocytes), (Neuroblast → Granule immature). We set the dimensions of **z**_*r*_ **h** to be 3 and 4 on this dataset.

#### Intestinal organoid

Data from [28], downloaded from dynamo [5]. The transitions tested with CBDir are (Stem cells → TA cells), (Stem cells → Goblet cells), (Stem cells → Tuft cells), (TA cells → Enterocytes). We set the dimensions of **z**_*r*_ and **h** to be 2 and 3 on this dataset. We use the celltype annotated model for this dataset.

#### Mouse hindbrain (Oligo)

Data of mouse hindbrain oligodendrocyte lineage from [10], downloaded from http://pklab.med.harvard.edu/ruslan/velocity/oligos/. The transitions tested with CBDir are (COPs → NFOLs), (NFOLs → MFOLs). We set the dimension of **z**_*r*_ and **h** to be 1 and 2 on this dataset. We use the celltype annotated model for this dataset.

#### Mouse hindbrain (GABA, Glial)

Processed data of mouse hindbrain with the differentiation of GABAergic interneuron and glial cells is downloaded from DeepVelo [18]. The transitions tested with CBDir are (Neural stem cells → progenitors), (Proliferating VZ progenitors → VZ progenitors), (VZ progenitors → Gliogenic progenitors), (VZ progenitors → Differentiating GABA interneurons), (Differentiating GABA interneurons → GABA interneurons). We set the dimensions of **z**_*r*_ and **h** to be 1 and 2 on this dataset. We use the celltype annotated model for this dataset.

#### Mouse erythroid

Erythroid lineage of mouse gastrulation [36]. Downloaded from the scVelo package [11]. The transitions tested with CBDir are (Blood progenitors 1 → Blood progenitors 2), (Blood progenitors 2 → Erythroid1), (Erythroid1 → Erythroid2), (Erythroid2 → Erythroid3). We set the dimension of **z**_*r*_ and **h** to be 1 and 1 on this dataset.

Human bone marrow. Data from [37], downloaded with scVelo [11]. The transitions tested with CBDir are (HSC 1 → CLP), (HSC 1 → Mega), (HSC 1 → Ery 1), (Ery 1 → Ery 2), (HSC 1 → HSC 2), (HSC 2 → Precursors), (HSC 2 → Mono 2), (Mono 2 → Mono 1), (Precursors → DCs). We set the dimension of **z**_*r*_ and **h** to be 2 and 2 on this dataset.

#### scNT-seq neuron KCl stimulation

Cortical neurons are stimulated with potassium chloride (KCl) for 0, 15, 30, and 60 minutes. Data from [46], downloaded from https://github.com/wulabupenn/scNT-seq. The transitions tested with CBDir are the times (0 → 15), (15 → 30), (30 → 60), (60 → 120). We set the dimension of **z**_*r*_ and **h** to be 1 and 1 on this dataset. We also restrict to only including the velocity genes in the likelihood.

#### Mouse embryonic fibroblast reprogramming

Reprogramming of mouse embryonic fibroblasts into induced endoderm progenitor cells [29]. Data was downloaded with the CellRank package [4]. We set the dimension of **z**_*r*_ to be 1 on this dataset. We use the celltype annotated model on this dataset.

#### Mouse gastrulation

Mouse gastrulation including all progenitors of major organs [36]. Downloaded from the scVelo package [11]. We subset to 20000 cells selected randomly for faster training. We set the dimensions of **z**_*r*_ and **h** to be 6 and 7 on this dataset. We use the celltype annotated model for this dataset.

#### Embryonic mouse brain

Data downloaded from 10X https://www.10xgenomics.com/resources/datasets/fresh-embryonic-e-18-mouse-brain-5-k-1-standard-1-0-0. We followed the pre-processing from MultiVelo [20]. We set the dimension of **z**_*r*_ to be 2 on this dataset.

## L. Code availability

LatentVelo is available at https://github.com/Spencerfar/LatentVelo. Code reproducing the results of the paper is also given.

## VI. ACKNOWLEDGEMENTS

We thank Eric Johnson and Dominic Skinner for reading and providing feedback on the manuscript. SF received funding from a University of Toronto Data Sciences Institute Posterdoctoral fellowship. The authors received funding from a University of Toronto Medicine by Design grant.

## SUPPLEMENTAL FIGURES

**FIG. S1.**
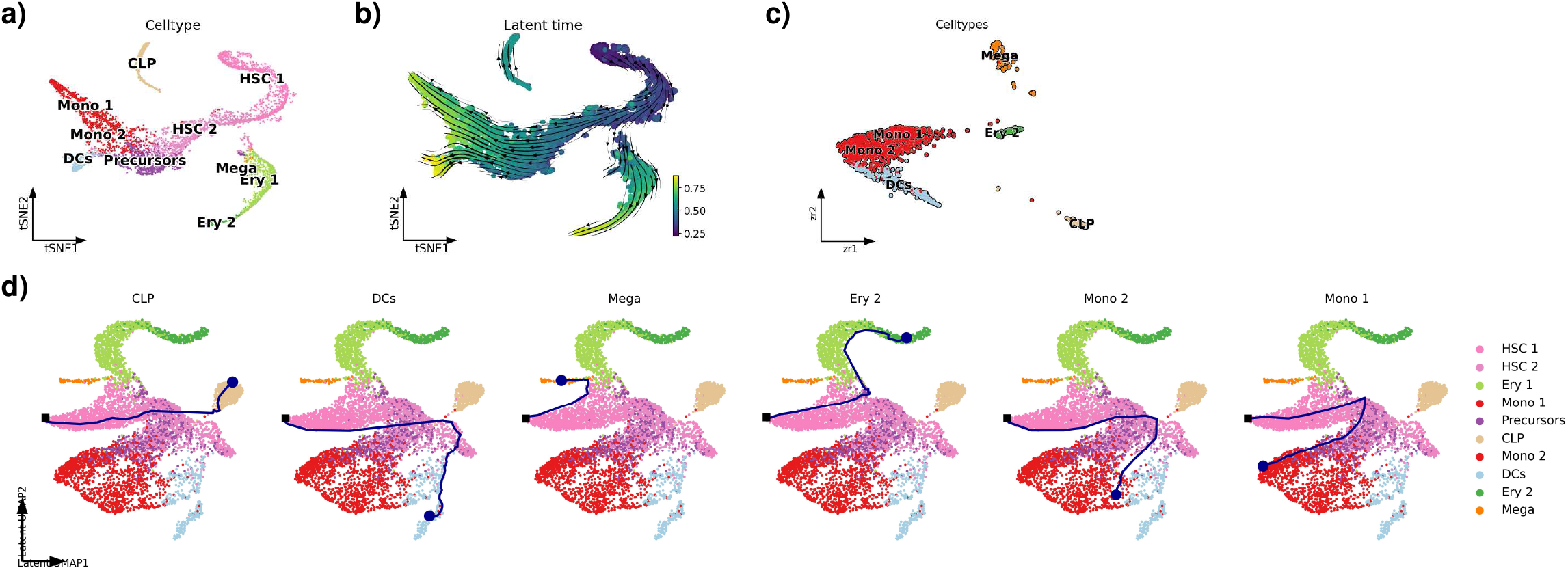
LatentVelo infers latent cell dynamics in human bonemarrow. **a)** tSNE plot showing the differentiation of hematopoietic stem cells into erythroid, common lymphocyte progenitor, monocyte, dendridic, and megakaryocyte cells. **b)** We show the velocity and latent time inferred by LatentVelo, indicating the directions of differentiation. **c)** We also show the inferred latent regulatory states in LatentVelo, highlighting the distinct dynamics for the terminal states. **d)** We show the latent trajectories inferred by LatentVelo for 6 different cells, corresponding to a CLP, DC, Mega, Ery, or Mono cell. Plot is shown for a UMAP representation of the spliced latent state. Dynamics start from the initial state **z**_0_ at the black square (the learned initial state) and follow the magenta path until terminating at the magenta circles at the estimated 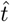. These trajectories indicate transitions that are initially unclear from the original gene-space tSNE plot in **a)**. In **a)**, the CLP cells are disconnected from the rest, but the shown trajectory indicates a transition from HSC cells. In **a)**, it is unclear whether the “precursor” cells are an intermediate celltype before DC cells, whereas the trajectory rather suggests that they are an intermediate celltype before Mono cells. In **a)**, it is unclear whether the Mono 1 and 2 cells are distinct Monocytes, or whether Mono 2 is an intermediate celltype before Mono 1, whereas the trajectories suggest that these are two distinct types of monocytes.

**FIG. S2.**
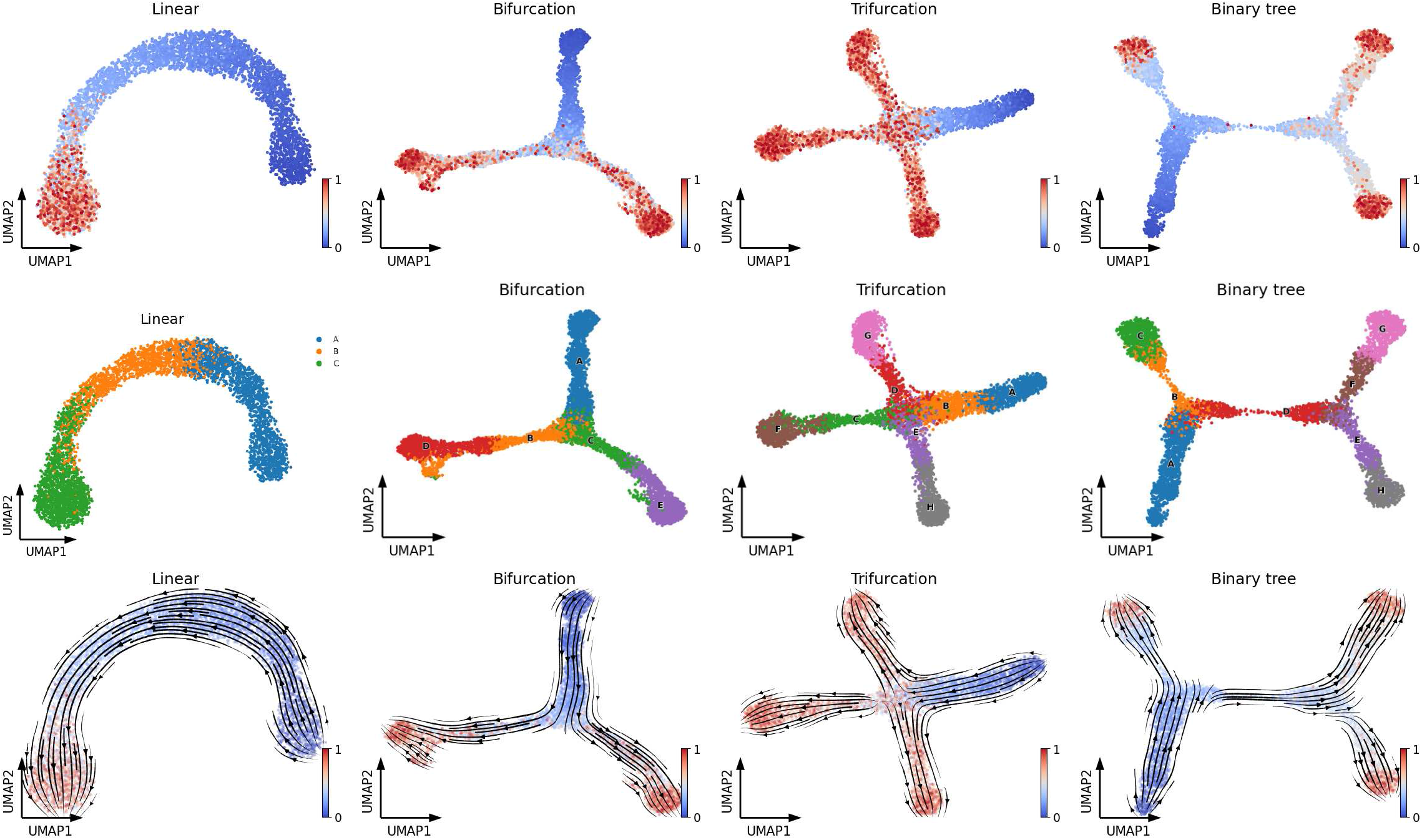
Synthetic datasets. UMAP plots showing the synthetic datasets used to benchmark models. Colored by simulation time and milestone, which is used for CBDir metric and the celltype annotated model. We also show velocity estimated by LatentVelo, colored by LatentVelo latent time.

**FIG. S3.**
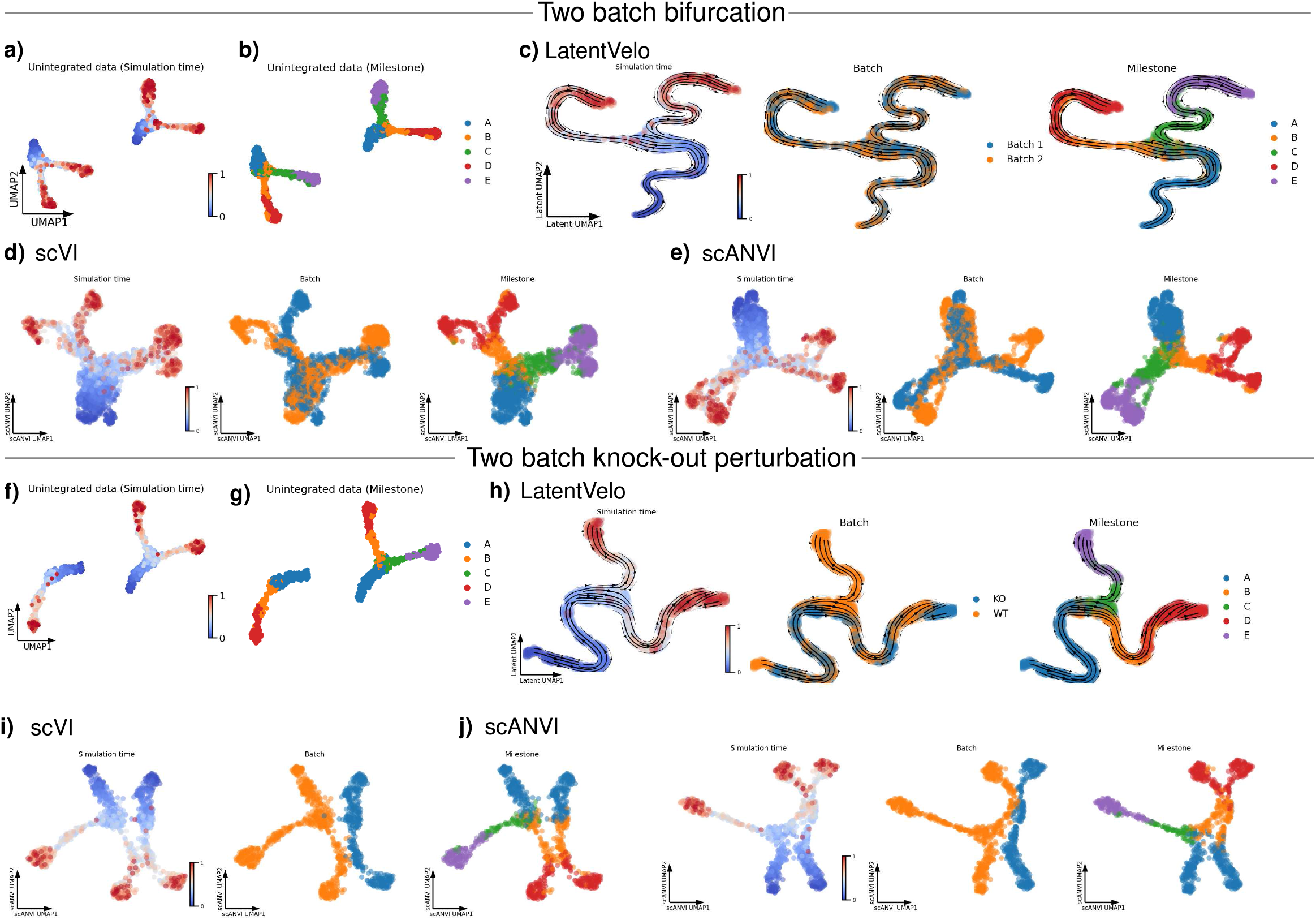
Comparison of batch correction for LatentVelo and scVI and scANVI. UMAP plots showing batch correction of synthetic datasets using LatentVelo, scVI [30], and scANVI [15]. **a)** and **b)** UMAP plots of the raw unintegrated batches for 2 batches of a bifurcation, colored by simulation time and milestone cells, highlighting the direction of differentiation and branches. **c)** UMAP plot of the LatentVelo latent space for this dataset. Velocity is in the correction direction, and clusters are well integrated. **d)** scVI and **e)** scANVI latent space UMAP plots. We see that these methods do not perform as well as LatentVelo. **f)** and **g)** UMAP plots of the raw unintegrated batches for 2 batches of a bifurcation with a gene module knockout, causing one branch to be eliminated. **h)** UMAP plot of the LatentVelo latent space for this dataset. Velocity is in the correction direction, and clusters are well integrated. **i)** scVI and **j)** scANVI latent space UMAP plots. We see that these methods fail to integrate these batches well.

**FIG. S4.**
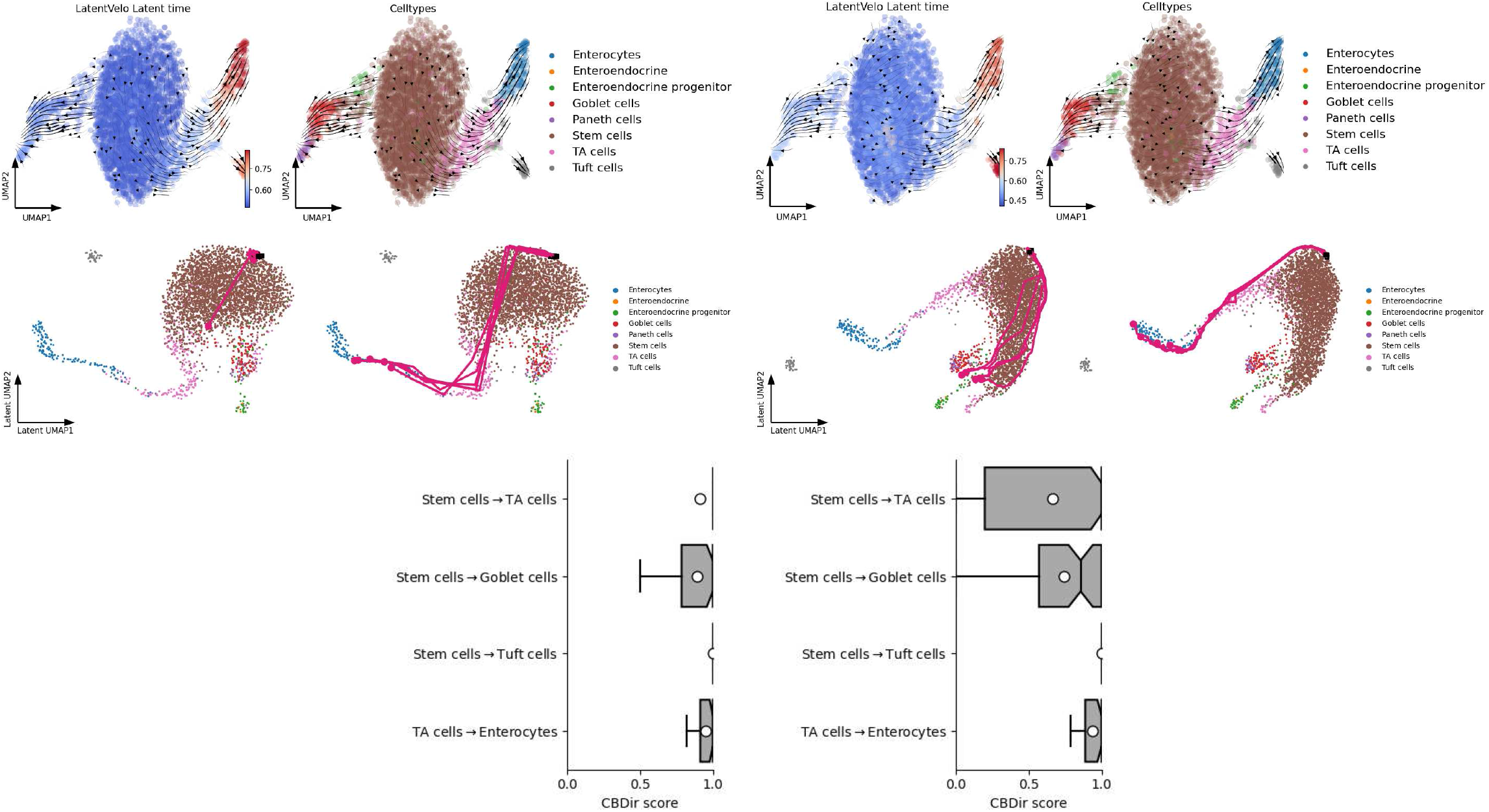
Exploring the role of latent space dimensionality. We use an intestional organoid dataset [27] to explore the dimensionality of the latent space in LatentVelo. (Left) we show a latent space dimension of 20, with the dimension of *z*_*r*_ as 2, and (right) with a latent space dimension of 75 and the dimension of *z*_*r*_ is also 2. Even though the velocities projected on the gene-space UMAP plot largely agree with the expected directions in both models, the underlying dynamics and inferred trajectories are distinct. We sample 5 trajectories for each of the branches in the bifurcation: trajectories for cells belonging to the enterocyte branch and cells belonging to the goblet/paneth cell branch. On the enterocyte branch, both models infer along the branch. However, the 20-dimensional model poorly infers trajectories for the goblet/paneth cell branch, with most trajectories staying near the **z**_0_.

**FIG. S5.**
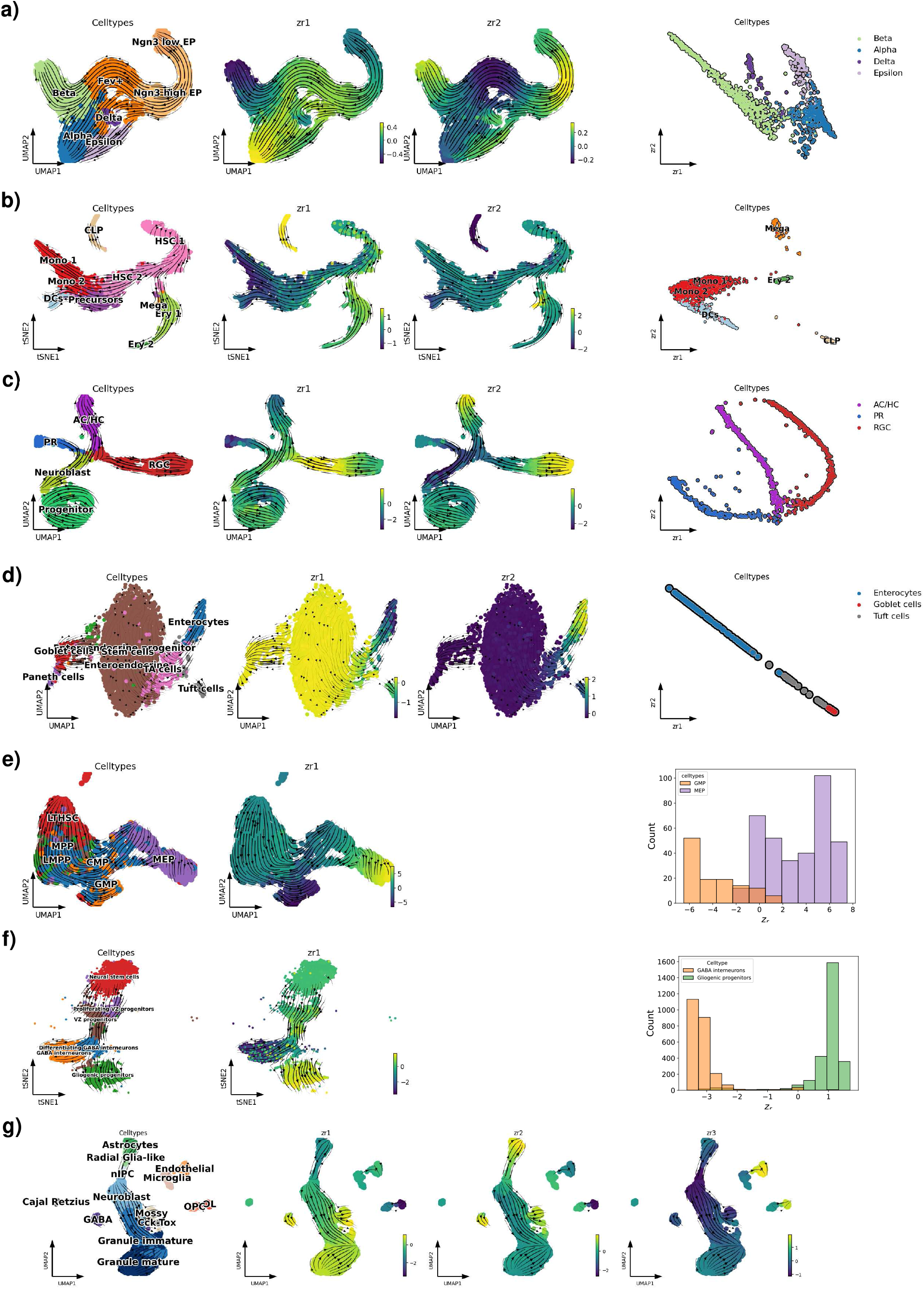
LatentVelo regulatory parameter z_*r*_ infers lineages. **a)** Pancreas, **b)** Human bone marrow, **c)** Retina, **d)** Intestinal organoid, **e)** Mouse hematopoiesis, **f)** hindbrain (GABA, glial), and **g)** dentate gyrus UMAP or tSNE plots shown with velocity arrows and colored by cell-types and regulatory parameters **z**_*r*_. In the case of 2 regulatory parameters, we show a scatter plot. For 1 regulatory parameter, we show a histogram. These plots show clear separation between the lineages, identified by **z**_r_. We also use a logistic regression classifier to predict terminal states from these low-dimension **z**_r_, achieving mean 25-fold cross-validation of accuracy of above 90% for all datasets, demonstrating that these clearly separated cell-type clusters accurately represent the different lineages. Since the latent dynamics depend on **z**_r_, these plots demonstrate the distinct dynamics on different lineages.

**FIG. S6.**
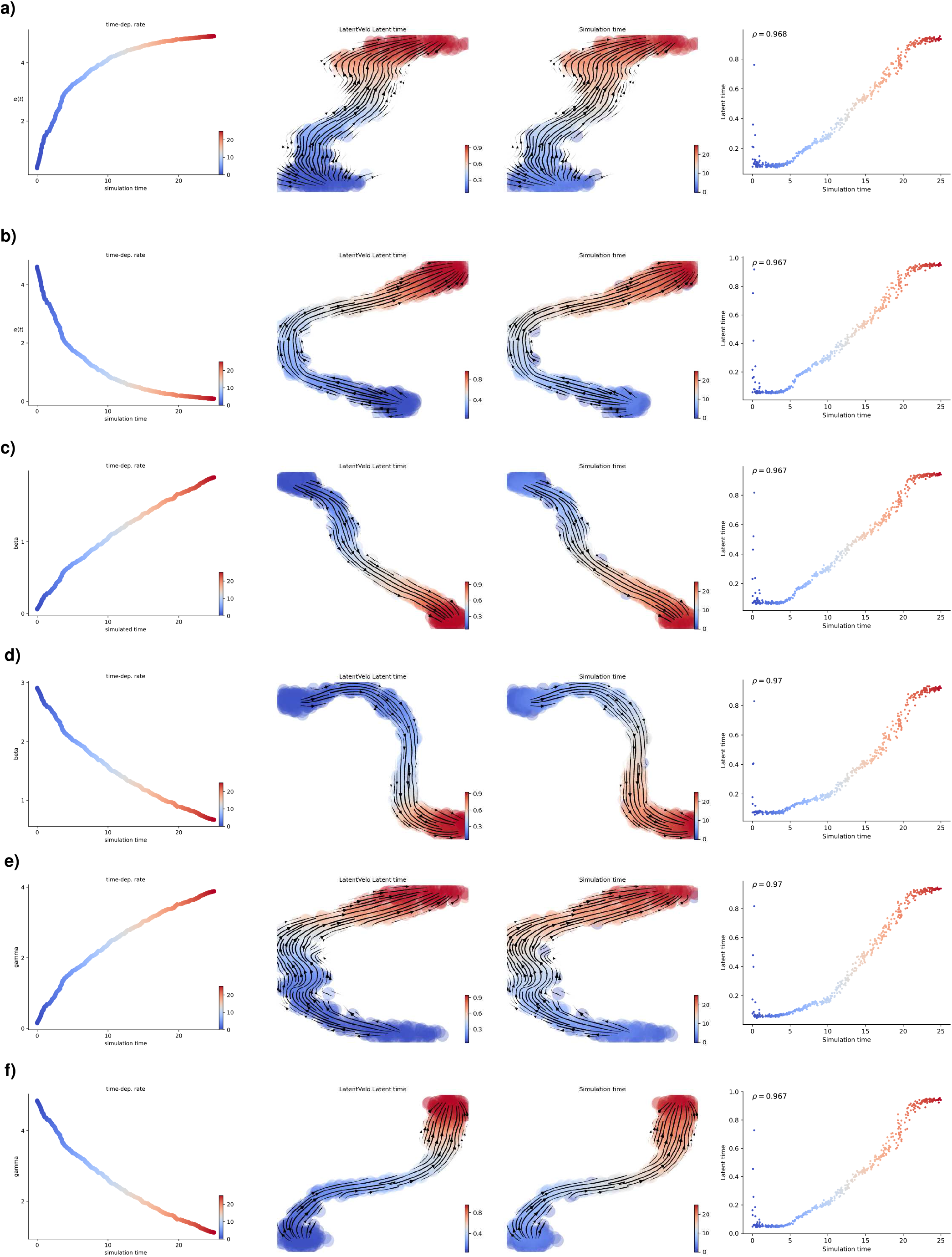
Time-dependent rate simulations. We simulate from the scVelo linear ODE model [11] while varying alpha (transcription rate), beta (splicing rate), and gamma (degradation rate). The simulation is run with 30 genes, and 5 are selected to be time-varying. We show increasing/decreasing for each of *α* (**a)** and **b)**), *β* (**c)** and **d)**), and *γ* (**e)** and **f)**). We assess our LatentVelo’s ability to infer the proper direction of differentiation by rank correlation between latent time and simulation time. We use LatentVelo with 10 latent dimensions for these small datasets.

## SUPPLEMENTAL INFORMATION

### 1. Model estimates error in latent time and velocity estimates

**FIG. S7.**
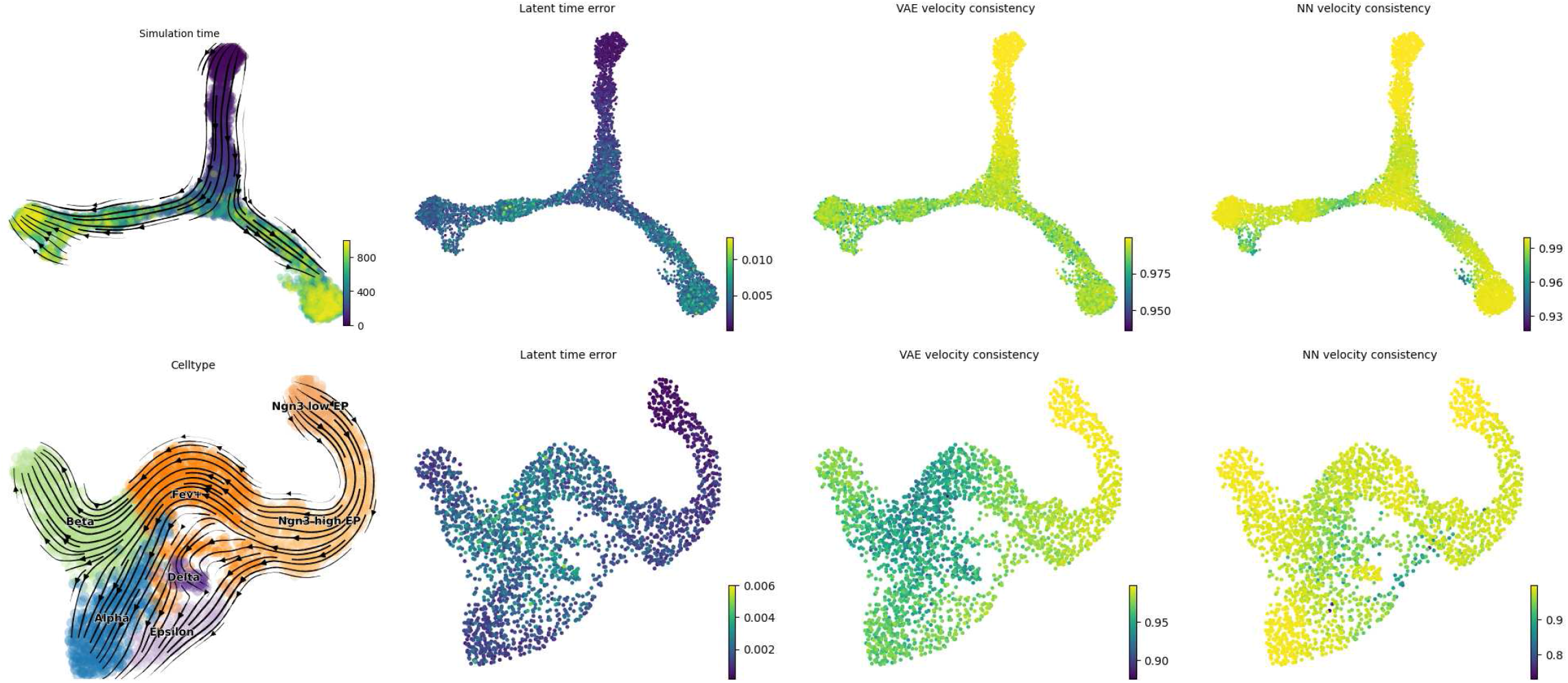
Error estimates in latent time and RNA velocity. (Top) Synthetic bifurcation. (Bottom) Pancreas dataset. Samples from the approximate Bayesian posterior distribution of latent time and RNA velocity are used to estimate the error (standard error of the mean) of latent time and the consistency of RNA velocity. as a comparison, we show the average nearest neighbor velocity consistency, introduced in the scVelo package [11].

Previous models of RNA velocity do not include uncertainty estimates in parameters. Our model infers the approximate Bayesian posterior distribution of the latent time and velocities, enabling the estimation of approximate uncertainties. However, we note that variational Bayesian approximations underestimate uncertainty [47], and so here we only interpret these uncertainty estimates to be *relative* estimates of uncertainty.

In Figure S7, we show the velocity inferred by the model on a UMAP plot, as well as the standard error of the mean of the latent time, the velocity consistency by re-sampling from the VAE latent space, and the consistency of velocity from nearest neighbors (as computed by scVelo [11]). The VAE velocity consistency is computed by the average cosine similarity of all pairs of samples from the VAE.

For the synthetic bifurcation dataset, we show that error increases near and after the bifurcation. Similarly with the pancreas dataset, where error increases in the area near the transitions to Alpha, Beta, Delta, and Epsilon cells, and then decreases in the Alpha and Beta cells after the transition is made. These estimates make sense, showing a decreased uncertainty where cell lineage decisions are being made.

### 2. Integrating multi-omic data with ATAC-seq

To show our approach can be generalized to other data modalities with multiomics, we use a mouse embryonic brain dataset incorporating both RNA-seq and ATAC-seq, previously used with MultiVelo [20]. The diagram in Figure S8 shows the new structure of the latent variables, where we now include the latent space representation of chromatin accessibility **z**_*c*_. In this model, chromatin accessibility effects transcription of unspliced RNA **z**_*u*_, and chromatin dynamics are regulated by **z**_*r*_. We still also allow the direct regulation of transcription by other methods than just chromatin accessibility, shown by retaining the connection between **z**_*u*_ and **z**_*r*_ as in Figure 1 above. The form of the dynamics for this model are:

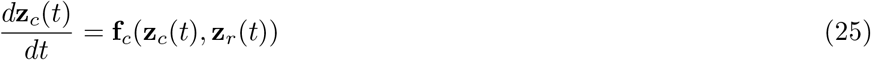

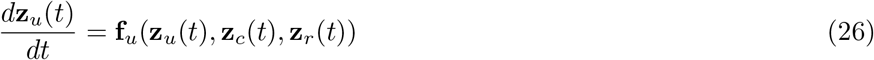

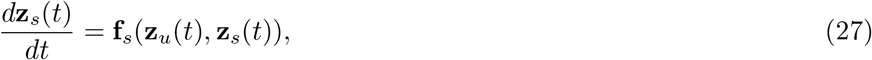

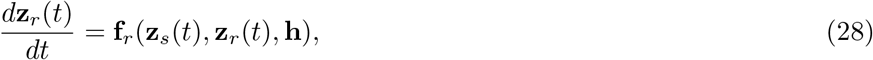

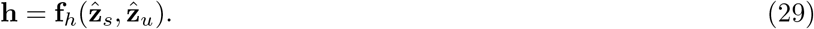

In addition to these equations, we adapt the correlation regularization on gene-space for the unspliced dynamics, and now use the regularization,

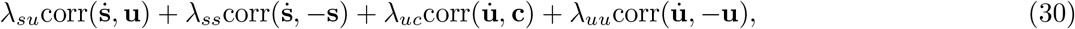

to enforce the direction of transition based on chromatin accessibility in addition to splicing. We use the same default *λ*_*su*_ = *λ*_*ss*_ = *λ*_*uc*_ = *λ*_*uu*_ = 0.1

In Figure S8 the RNA velocity on the UMAP plot shows the differentiation direction from radial glia cells to neurons in the main branch, and the differentiation of radial glia cells to interneurons on the other branches. The spliced RNA, unspliced RNA, and chromatin accessibility plots vs latent time for 4 different genes show the spliced, unspliced, and chromatin velocities. Velocities are positive when these variables increase, and negative when they decrease, as expected.

**FIG. S8.**
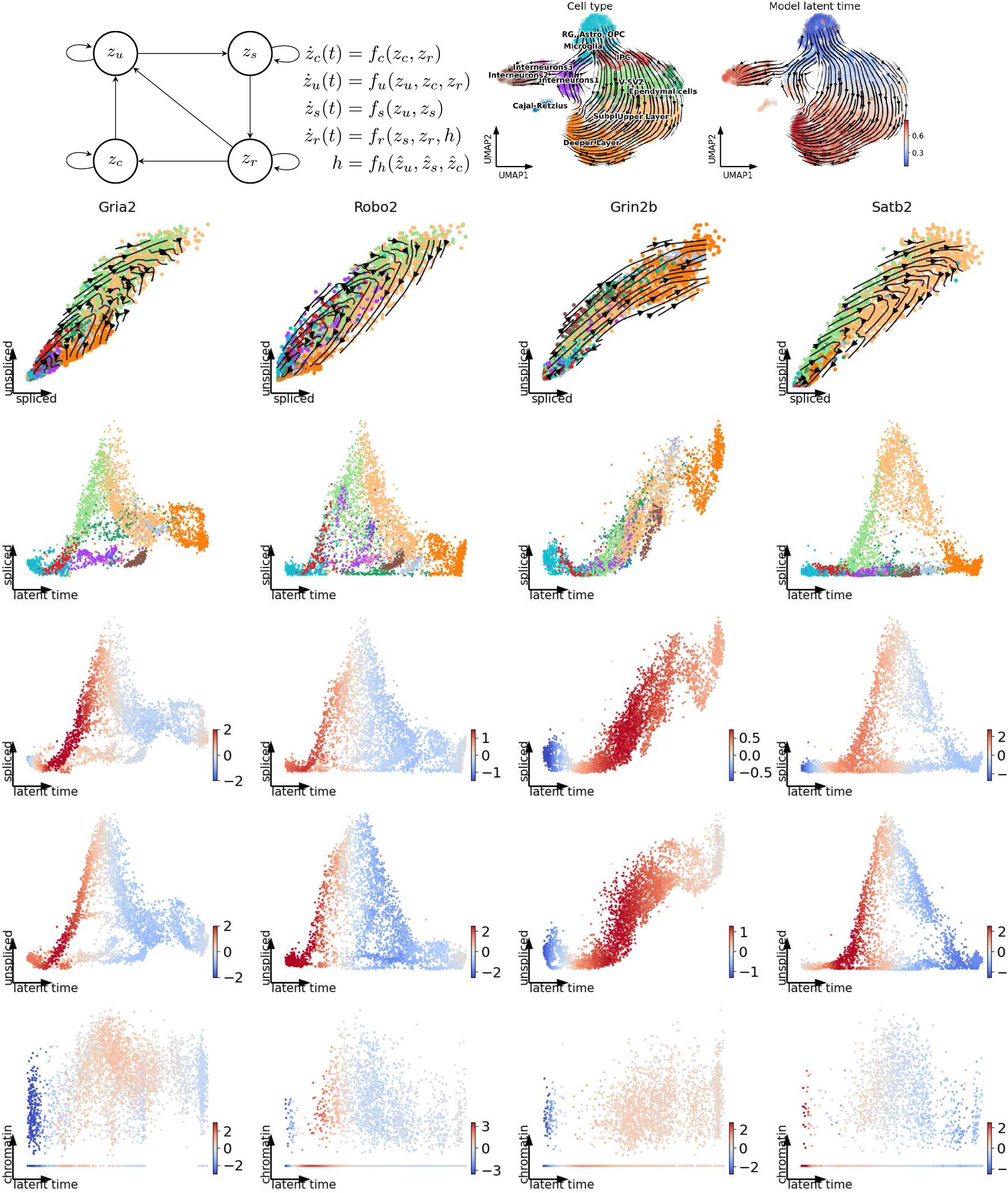
Integration of chromatin accessibility. **a)** Integration of chromatin accessibility into the structured latent dynamics. We incorporate this into latentVelo by allowing the latent representation of chromatin accessibility *z*_c_ to influence the transcription of unspliced RNA *z*_*u*_. We also include the regulation of chromatin dynamics by the latent regularization variable *z*_*r*_. **b)** UMAP representation of a mouse embryonic brain dataset showing cell types and the inferred latent time and velocity with LatentVelo. **c)** Spliced RNA, unspliced RNA, and chromatin accessibility for 4 selected genes, colored by the corresponding chromatin, unspliced, and spliced velocity.

## References

[1] Louise Deconinck, Robrecht Cannoodt, Wouter Saelens, Bart Deplancke, and Yvan Saeys. Recent advances in trajectory inference from single-cell omics data. Current Opinion in Systems Biology, 27:100344, September 2021.

[2] Wouter Saelens, Robrecht Cannoodt, Helena Todorov, and Yvan Saeys. A comparison of single-cell trajectory inference methods. Nat. Biotechnol., 37(5):547–554, May 2019.

[3] Jun Ding, Nadav Sharon, and Ziv Bar-Joseph. Temporal modelling using single-cell transcriptomics. Nat. Rev. Genet., 23(6):355–368, June 2022.

[4] Marius Lange, Volker Bergen, Michal Klein, Manu Setty, Bernhard Reuter, Mostafa Bakhti, Heiko Lickert, Meshal Ansari, Janine Schniering, Herbert B Schiller, Dana Pe’er, and Fabian J Theis. CellRank for directed single-cell fate mapping. Nat. Methods, 19(2):159–170, January 2022.

[5] Xiaojie Qiu, Yan Zhang, Jorge D Martin-Rufino, Chen Weng, Shayan Hosseinzadeh, Dian Yang, Angela N Pogson, Marco Y Hein, Kyung Hoi Joseph Min, Li Wang, Emanuelle I Grody, Matthew J Shurtleff, Ruoshi Yuan, Song Xu, Yian Ma, Joseph M Replogle, Eric S Lander, Spyros Darmanis, Ivet Bahar, Vijay G Sankaran, Jianhua Xing, and Jonathan S Weissman. Mapping transcriptomic vector fields of single cells. Cell, 185(4):690–711.e45, February 2022.

[6] Ziqi Zhang and Xiuwei Zhang. Inference of high-resolution trajectories in single-cell RNA-seq data by using RNA velocity. Cell Reports Methods, 1(6):100095, October 2021.

[7] R Gupta, D Cerletti, G Gut, A Oxenius, and M Claassen. Cytopath: Simulation based inference of differentiation trajectories from RNA velocity fields. August 2021.

[8] Zhanlin Chen, William C King, Aheyon Hwang, Mark Gerstein, and Jing Zhang. DeepVelo: Single-cell transcriptomic deep velocity field learning with neural ordinary differential equations. April 2022.

[9] Ruishan Liu, Angela Oliveira Pisco, Emelie Braun, Sten Linnarsson, and James Zou. Dynamical systems model of RNA velocity improves inference of single-cell trajectory, pseudo-time and gene regulation. J. Mol. Biol., page 167606, April 2022.

[10] Gioele La Manno, Ruslan Soldatov, Amit Zeisel, Emelie Braun, Hannah Hochgerner, Viktor Petukhov, Katja Lidschreiber, Maria E Kastriti, Peter Lönnerberg, Alessandro Furlan, Jean Fan, Lars E Borm, Zehua Liu, David van Bruggen, Jimin Guo, Xiaoling He, Roger Barker, Erik Sundström, Gonçalo Castelo-Branco, Patrick Cramer, Igor Adameyko, Sten Linnarsson, and Peter V Kharchenko. RNA velocity of single cells. Nature, 560(7719):494–498, August 2018.

[11] Volker Bergen, Marius Lange, Stefan Peidli, F Alexander Wolf, and Fabian J Theis. Generalizing RNA velocity to transient cell states through dynamical modeling. Nat. Biotechnol., 38(12):1408–1414, August 2020.

[12] Melania Barile, Ivan Imaz-Rosshandler, Isabella Inzani, Shila Ghazanfar, Jennifer Nichols, John C Marioni, Carolina Guibentif, and Berthold Göttgens. Coordinated changes in gene expression kinetics underlie both mouse and human erythroid maturation. Genome Biol., 22(1):197, July 2021.

[13] Volker Bergen, Ruslan A Soldatov, Peter V Kharchenko, and Fabian J Theis. RNA velocity—current challenges and future perspectives. Mol. Syst. Biol., 17(8):e10282, August 2021.

[14] Gennady Gorin, Meichen Fang, Tara Chari, and Lior Pachter. RNA velocity unraveled. February 2022.

[15] Chenling Xu, Romain Lopez, Edouard Mehlman, Jeffrey Regier, Michael I Jordan, and Nir Yosef. Probabilistic harmonization and annotation of single-cell transcriptomics data with deep generative models. Mol. Syst. Biol., 17(1):e9620, January 2021.

[16] Chen Qiao and Yuanhua Huang. Representation learning of RNA velocity reveals robust cell transitions. Proceedings of the National Academy of Sciences, 118(49):e2105859118, 2021.

[17] Mingze Gao, Chen Qiao, and Yuanhua Huang. UniTVelo: temporally unified RNA velocity reinforces single-cell trajectory inference. April 2022.

[18] Haotian Cui, Hassaan Maan, and Bo Wang. DeepVelo: Deep learning extends RNA velocity to multi-lineage systems with cell-specific kinetics. April 2022.

[19] Yichen Gu, David Blaauw, and Joshua D. Welch. Bayesian inference of rna velocity from multi-lineage single-cell data. bioRxiv, 2022.

[20] Chen Li, Maria Virgilio, Kathleen L Collins, and Joshua D Welch. Single-cell multi-omic velocity infers dynamic and decoupled gene regulation. December 2021.

[21] Yuanhua Huang and Guido Sanguinetti. BRIE2: computational identification of splicing phenotypes from single-cell transcriptomic experiments. Genome Biol., 22(1):251, August 2021.

[22] Qian Li. sctour: a deep learning architecture for robust inference and accurate prediction of cellular dynamics. April 2022.

[23] Diederik P. Kingma and Max Welling. Auto-encoding variational bayes. In Yoshua Bengio and Yann LeCun, editors, 2nd International Conference on Learning Representations, ICLR 2014, Banff, AB, Canada, April 14-16, 2014, Conference Track Proceedings, 2014.

[24] Danilo Jimenez Rezende, Shakir Mohamed, and Daan Wierstra. Stochastic backpropagation and approximate inference in deep generative models. In Proceedings of the 31st International Conference on International Conference on Machine Learning -Volume 32, ICML’14, pages II–1278–II–1286. JMLR.org, June 2014.

[25] Ricky T. Q. Chen, Yulia Rubanova, Jesse Bettencourt, and David K Duvenaud. Neural ordinary differential equations. In S. Bengio, H. Wallach, H. Larochelle, K. Grauman, N. Cesa-Bianchi, and R. Garnett, editors, Advances in Neural Information Processing Systems, volume 31. Curran Associates, Inc., 2018.

[26] Malte D Luecken, M Büttner, K Chaichoompu, A Danese, M Interlandi, M F Mueller, D C Strobl, L Zappia, M Dugas, M Colomé-Tatché, and Fabian J Theis. Benchmarking atlas-level data integration in single-cell genomics. Nat. Methods, 19(1):41–50, December 2021.

[27] Aimée Bastidas-Ponce, Sophie Tritschler, Leander Dony, Katharina Scheibner, Marta Tarquis-Medina, Ciro Salinno, Silvia Schirge, Ingo Burtscher, Anika Böttcher, Fabian J Theis, Heiko Lickert, and Mostafa Bakhti. Comprehensive single cell mRNA profiling reveals a detailed roadmap for pancreatic endocrinogenesis. Development, 146(12), June 2019.

[28] Nico Battich, Joep Beumer, Buys de Barbanson, Lenno Krenning, Chloé S Baron, Marvin E Tanenbaum, Hans Clevers, and Alexander van Oudenaarden. Sequencing metabolically labeled transcripts in single cells reveals mRNA turnover strategies. Science, 367(6482):1151–1156, 2020.

[29] Brent A Biddy, Wenjun Kong, Kenji Kamimoto, Chuner Guo, Sarah E Waye, Tao Sun, and Samantha A Morris. Single-cell mapping of lineage and identity in direct reprogramming. Nature, 564(7735):219–224, December 2018.

[30] Romain Lopez, Jeffrey Regier, Michael B Cole, Michael I Jordan, and Nir Yosef. Deep generative modeling for single-cell transcriptomics. Nat. Methods, 15(12):1053–1058, December 2018.

[31] Robrecht Cannoodt, Wouter Saelens, Louise Deconinck, and Yvan Saeys. Spearheading future omics analyses using dyngen, a multi-modal simulator of single cells. Nat. Commun., 12(1):1–9, June 2021.

[32] Jolene S Ranek, Natalie Stanley, and Jeremy E Purvis. Integrating temporal single-cell gene expression modalities for trajectory inference and disease prediction. March 2022.

[33] Kasper Daniel Hansen. Batch effects in scrna velocity analysis, 2021.

[34] W Evan Johnson, Cheng Li, and Ariel Rabinovic. Adjusting batch effects in microarray expression data using empirical bayes methods. Biostatistics, 8(1):118–127, April 2006.

[35] Mohammad Lotfollahi, F Alexander Wolf, and Fabian J Theis. scgen predicts single-cell perturbation responses. Nat. Methods, 16(8):715–721, July 2019.

[36] Blanca Pijuan-Sala, Jonathan A Griffiths, Carolina Guibentif, Tom W Hiscock, Wajid Jawaid, Fernando J Calero-Nieto, Carla Mulas, Ximena Ibarra-Soria, Richard C V Tyser, Debbie Lee Lian Ho, Wolf Reik, Shankar Srinivas, Benjamin D Simons, Jennifer Nichols, John C Marioni, and Berthold Göttgens. A single-cell molecular map of mouse gastrulation and early organogenesis. Nature, 566(7745):490–495, February 2019.

[37] Manu Setty, Vaidotas Kiseliovas, Jacob Levine, Adam Gayoso, Linas Mazutis, and Dana Pe’er. Characterization of cell fate probabilities in single-cell data with palantir. Nat. Biotechnol., 37(4):451–460, March 2019.

[38] Adam Gayoso, Philipp Weiler, Mohammad Lotfollahi, Dominik Klein, Justin Hong, Aaron Streets, Fabian J. Theis, and Nir Yosef. Deep generative modeling of transcriptional dynamics for rna velocity analysis in single cells. bioRxiv, 2022.

[39] Qian Qin, Eli Bingham, Gioele La Manno, David M. Langenau, and Luca Pinello. Pyro-velocity: Probabilistic rna velocity inference from single-cell data. bioRxiv, 2022.

[40] Xuechen Li, Ting-Kam Leonard Wong, Ricky T. Q. Chen, and David Duvenaud. Scalable gradients for stochastic differential equations. International Conference on Artificial Intelligence and Statistics, 2020.

[41] Grace Hui Ting Yeo, Sachit D Saksena, and David K Gifford. Generative modeling of single-cell time series with PRE-SCIENT enables prediction of cell trajectories with interventions. Nat. Commun., 12(1):1–12, May 2021.

[42] Ricky T. Q. Chen. torchdiffeq, 2018.

[43] F Alexander Wolf, Philipp Angerer, and Fabian J Theis. SCANPY: large-scale single-cell gene expression data analysis. Genome Biol., 19(1):15, February 2018.

[44] Adam Gayoso, Romain Lopez, Galen Xing, Pierre Boyeau, Valeh Valiollah Pour Amiri, Justin Hong, Katherine Wu, Michael Jayasuriya, Edouard Mehlman, Maxime Langevin, Yining Liu, Jules Samaran, Gabriel Misrachi, Achille Nazaret, Oscar Clivio, Chenling Xu, Tal Ashuach, Mariano Gabitto, Mohammad Lotfollahi, Valentine Svensson, Eduardo da Veiga Bel-trame, Vitalii Kleshchevnikov, Carlos Talavera-López, Lior Pachter, Fabian J. Theis, Aaron Streets, Michael I. Jordan, Jeffrey Regier, and Nir Yosef. A python library for probabilistic analysis of single-cell omics data. Nature Biotechnology, Feb 2022.

[45] Sonia Nestorowa, Fiona K Hamey, Blanca Pijuan Sala, Evangelia Diamanti, Mairi Shepherd, Elisa Laurenti, Nicola K Wilson, David G Kent, and Berthold Göttgens. A single-cell resolution map of mouse hematopoietic stem and progenitor cell differentiation. Blood, 128(8):e20–e31, August 2016.

[46] Qi Qiu, Peng Hu, Xiaojie Qiu, Kiya W Govek, Pablo G Cámara, and Hao Wu. Massively parallel and time-resolved rna sequencing in single cells with scnt-seq. Nature methods, 17(10):991–1001, 2020.

[47] David M. Blei, Alp Kucukelbir, and Jon D. McAuliffe. Variational inference: A review for statisticians. Journal of the American Statistical Association, 112(518):859–877, 2017.

